# A Drug–Target Specificity Foundation Model for Off-target Prediction, Repurposing, and Generative Design

**DOI:** 10.64898/2026.06.08.730844

**Authors:** Sai T. Reddy

## Abstract

Molecular recognition - which small molecule binds which protein, and with what selectivity - governs the efficacy, safety, and discovery of every therapeutic, yet binding specificity is still determined by experimental screening or by computational methods that first predict three-dimensional structure. Transformer softmax attention is mathematically isomorphic to the Boltzmann distribution governing molecular binding at thermal equilibrium^1^, an identity that prescribes a single sequence-native architecture: the Specificity Foundation Model (SFM), which computes molecular binding compatibility as a thermodynamic quantity directly from sequence^2^. The framework was recently realized as prototype encoders across six molecular-recognition domains^3^. Here we report the small molecule drug-target protein SFM (dtSFM) as the first instance to pair a full-scale encoder with a generative decoder, trained on publicly available data consisting of 714,747 measured drug–protein interactions spanning 522,776 compounds and 22,964 proteins. Throughout, we verify binding predictions with AlphaFold 3^4^ as an orthogonal structural verifier that shares no architecture, training data, or representational basis with dtSFM. From this single dtSFM model we demonstrate the three sequence-native applications of drug discovery: off-target prediction, repurposing, and generative design. The dtSFM encoder retrieves a drug’s target, and a target’s drug, at 95% and 89% recall-at-10 in distribution, respectively. In the drug→target direction it screens off-targets at proteome scale, ranking the documented off-targets of clinical kinase inhibitors at a median of 30th out of 4,910 genes - the top 0.6% of the screen - when validated against a chemoproteomic panel^5^. In the target→drug direction it ranks the full 522,776-compound library against three immunology targets, identifying 46 novel candidates that pass AlphaFold-3 structural gating. The dtSFM cross-attentive decoder generates novel molecules for 16 targets, 850 of 1,200 (71%) designed candidates match the AlphaFold 3 structural confidence of the approved drug (iPTM ≥ 0.9 and interface PAE ≤ 1.67 Å), with the best candidates reaching iPTM 0.95–0.99 and interface PAE 0.79–1.37 Å. dtSFM brings computational thermodynamics to every stage where molecular recognition shapes drug discovery; experimental wet-lab validation is the immediate next step.

## INTRODUCTION

The central computation of pharmacology is molecular recognition: which small molecule binds which protein, and with what selectivity. The same question, asked in three directions, defines three of the field’s hardest problems: *efficacy* (does a drug engage its intended target), *safety* (what else does it bind), and *discovery* (what new chemistry binds a target of interest). In each case the determination is still made experimentally, by an industrialised pipeline that spans target identification, hit finding, hit-to-lead, and lead optimisation^6^. Off-target activity is mapped by chemoproteomic and kinase-panel screening with broad selectivity assays^5,7^; binding affinity is measured by surface plasmon resonance, isothermal titration calorimetry, biolayer interferometry, and orthogonal biophysical techniques^8^; and new starting points are advanced by high-throughput screening, fragment-based discovery, and medicinal-chemistry iteration in lead optimisation campaigns^9,10^.

The dominant computational paradigm approaches this through three-dimensional structure. AlphaFold 3 (AF3)^4^ and Boltz-1,-2 ^11,12^ predict the geometry of a drug–protein complex and report pose-confidence metrics that correlate with binding. Molecular design tools generate novel ligands by optimising structure directly: RFdiffusion^13^, BindCraft ^14^, and BoltzGen^15^ for protein binders; DiffSBDD^16^, TargetDiff ^17^, and Pocket2Mol ^18^ for small molecules. These methods are powerful and have transformed structural modelling, but they solve a different problem — predicting geometry — and benefit from or require structural information at inference.

We recently showed that the softmax attention mechanism of transformers^19^ is the Boltzmann distribution governing molecular binding at thermal equilibrium, with Luce’s uniqueness theorem^20^ establishing it as the only selection rule consistent with that physics^1^. From this identity and five conditions of molecular recognition systems, the architecture for any binding domain is prescribed — dual sequence encoders, symmetric contrastive training, and a learned temperature corresponding to physical *k_B_T* — defining a Specificity Foundation Model (SFM)^2^. An SFM does not approximate an unknown function; it estimates the parameters of a known one – the convergence equation. The probability that a drug binds a given target is the partition-function-normalised exponential of a learned compatibility score, so retrieval over a candidate pool is literally a thermodynamic selection computed from sequence. In this sense an SFM performs **computational thermodynamics**: it ranks binding by free-energy-like compatibility without ever constructing a structure. We have realised this framework first for antibody–antigen recognition^21^ and subsequently as prototype encoders across six molecular domains^3^. Here we train and test dtSFM, the first SFM realisation that combines a full-scale encoder, a generative decoder, and a deployable design loop in a single model.

dtSFM is trained on 714,747 paired drug–protein interactions covering 522,776 unique compounds and 22,964 unique proteins. The encoder achieves in-distribution recall-at-10 of 95% for drug → target retrieval (pool of 928 proteins, random baseline 1.1%) and 89% for target → drug retrieval (pool of 3,955 drugs, random baseline 0.25%). From this single model we report three applications. dtSFM performs off-target safety screening (drug → target) at proteome scale, recovering the kinome-wide off-target landscape of clinical kinase inhibitors against a published chemoproteomic off-target panel^5^ with mean AUROC 0.78. For library-scale repurposing (target → drug), dtSFM ranks the full 522,776-compound library against three clinically relevant targets (NLRP3, CD73, and STING1), identifying 46 novel candidates that pass structural confidence gating using AF3 ^4^ (iPTM ≥ 0.7 and interface PAE ≤ 5 Å). For generative design (target → novel drug), the first trained instance of the cross-attentive SFM decoder, applied to 16 clinically relevant targets, generates 1,200 novel molecules, of which 850 (71%) reach structural confidence matching that of the approved drug at each target (iPTM ≥ 0.9 and interface PAE ≤ 1.67 Å). Together these three applications demonstrate dtSFM as a single sequence-native foundation model spanning the molecular-recognition stages of computational drug discovery.

## RESULTS

### Constructing a drug–target specificity foundation model

Building on the antibody–antigen encoder CALM^21^ and the six prototype SFM encoders^3^, dtSFM is the first SFM realisation that combines a full-scale encoder, a generative decoder, and a deployable design loop in a single model. The architecture follows the convergence of the Boltzmann distribution for molecular binding with the softmax attention of transformers, which together with five conditions of molecular recognition systems prescribes the same dual-encoder + cross-attention + contrastive backbone for every binding domain^1,2^. dtSFM is the drug–target instantiation (**Fig. 1**). Two frozen pretrained encoders are joined by a trainable cross-attention encoder. The drug encoder is MoLFormer-XL^22^, which embeds a compound’s SMILES string^23^ into a 768-dimensional vector; the protein encoder is ESM-2-650M^24^, which embeds the target protein sequence into a 1,280-dimensional per-residue representation, attention-pooled over residues. Both encoders are frozen — their weights are not updated — and only the cross-attention encoder (two layers, eight heads, model dimension 512, feed-forward dimension 2,048; 14.4 million trainable parameters) and four output heads are trained. The asymmetric cross-attention fuses the two modalities into a joint drug–target representation from which the four heads read: (i) a global-retrieval head scoring a pair by cosine similarity and operating in both directions (drug→target for off-target safety screening, target→drug for repurposing), (ii) an interface head (per-atom binding-interface membership), (iii) a contact head (atom × residue contact map), (iv) and an affinity head (binding-strength regression).

**Figure 1.**
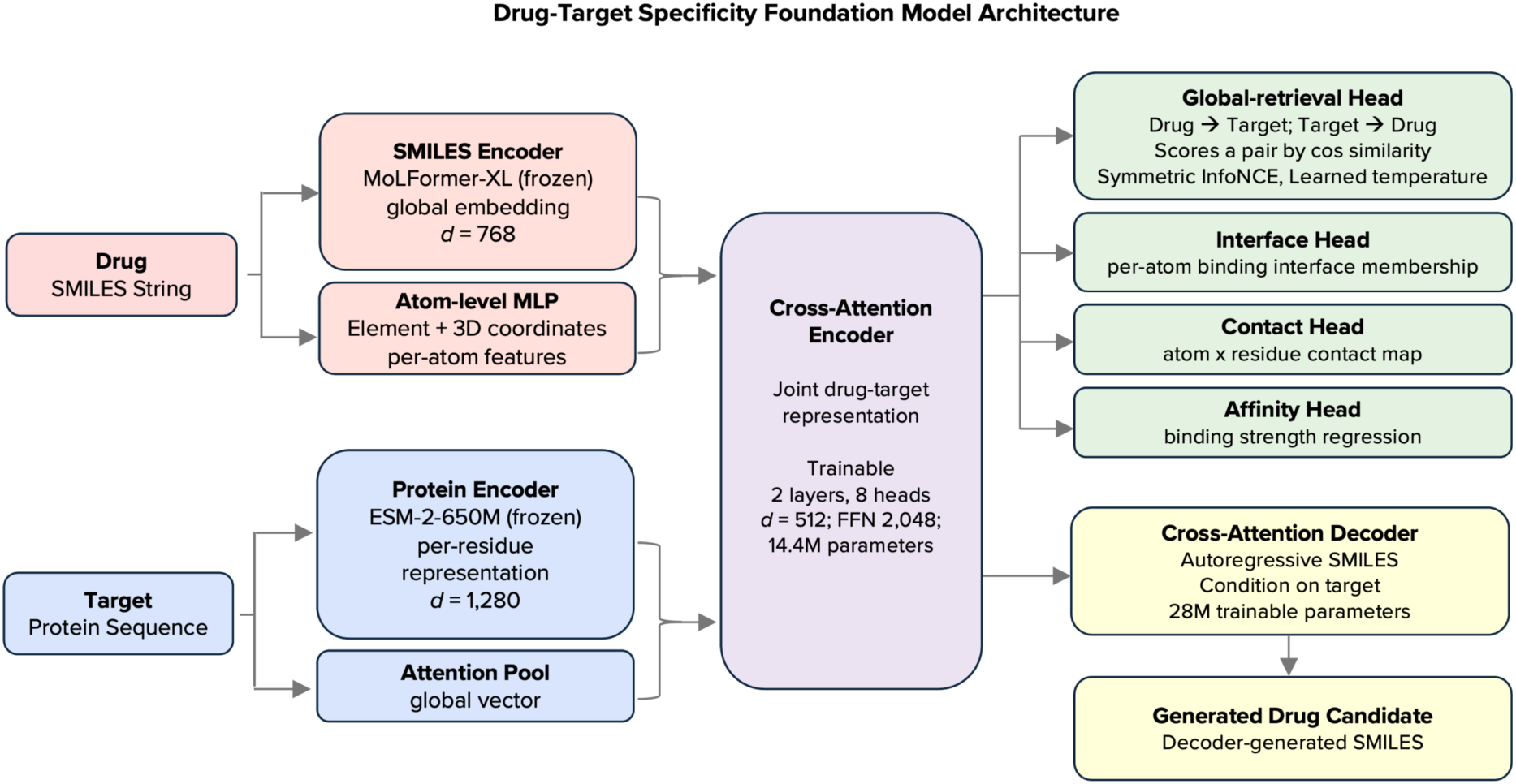
Architecture of the drug–target specificity foundation model (dtSFM). dtSFM fuses two frozen pretrained sequence encoders through a trainable cross-attention encoder and reads the joint representation with four task heads plus a generative decoder. The drug branch encodes the small-molecule SMILES with a frozen MoLFormer-XL transformer, contributing a 768→512 global embedding and per-atom features from an atom-level MLP over element identity and 3D coordinates; the protein branch encodes the target amino-acid sequence with a frozen ESM-2-650M transformer, contributing a 1,280→512 per-residue representation and an attention-pooled global vector. The trainable cross-attention encoder (2 layers, 8 heads, model dimension 512, feed-forward dimension 2,048; 14.4 M trainable parameters) fuses the two modalities into a joint drug–target representation. Four heads read this representation: a global-retrieval head scoring a pair by cosine similarity and operating bidirectionally (drug→target for off-target safety screening, target→drug for repurposing) under a symmetric InfoNCE loss with a learned temperature *t* = *kBT*; an interface head (per-atom binding-interface membership); a contact head (atom × residue contact map); and an affinity head (binding-strength regression, preliminary). A cross-attentive autoregressive decoder generates a novel SMILES string one token at a time, conditioned on the target via cross-attention over the per-residue features. Both input encoders are frozen; only the cross-attention encoder, heads, and decoder are trained.

The training corpus comprises 714,747 curated (drug, protein) pairs assembled from two sources (**Fig. 2a** and **Table 1**). PDBbind v2020 ^25^ provides 19,037 co-crystal complexes with experimentally measured *K_D_ / IC_50_*. The bulk of the corpus is drawn from the SAIR repository^26^, which provides 695,710 compound–protein pairs each combining a measured potency value with a Boltz-1x predicted binding interface retained at high prediction confidence (iPTM ≥ 0.70 and overall confidence score ≥ 0.70). The SAIR curation pipeline harvests bioactivities from the two largest public bioactivity databases — ChEMBL (release 33)^27^, which aggregates over 20 million curated bioactivity measurements from peer-reviewed publications and submitted datasets, and BindingDB^28^, which contains over 2.8 million binding affinities for more than 1.2 million compounds against more than 9,400 protein targets — and applies standardised assay-quality filters (binding-mode measurements, 1 pM to 100 μM dynamic range), ligand filters (PAINS exclusion, molecular weight ≤ 1,250 Da), and protein-quality filters (≤ 2,000 residues, monomeric assembly), while excluding any (compound, protein) system already represented in the PDB to prevent leakage against PDB-based training data. Each SAIR drug-target pair therefore couples an independently measured assay value with a high-confidence predicted complex — two orthogonal supports for the same interaction — which is why SAIR serves as primary supervision rather than synthetic augmentation. Together the two sources span 522,776 unique compounds and 22,964 unique proteins.

**Figure 2.**
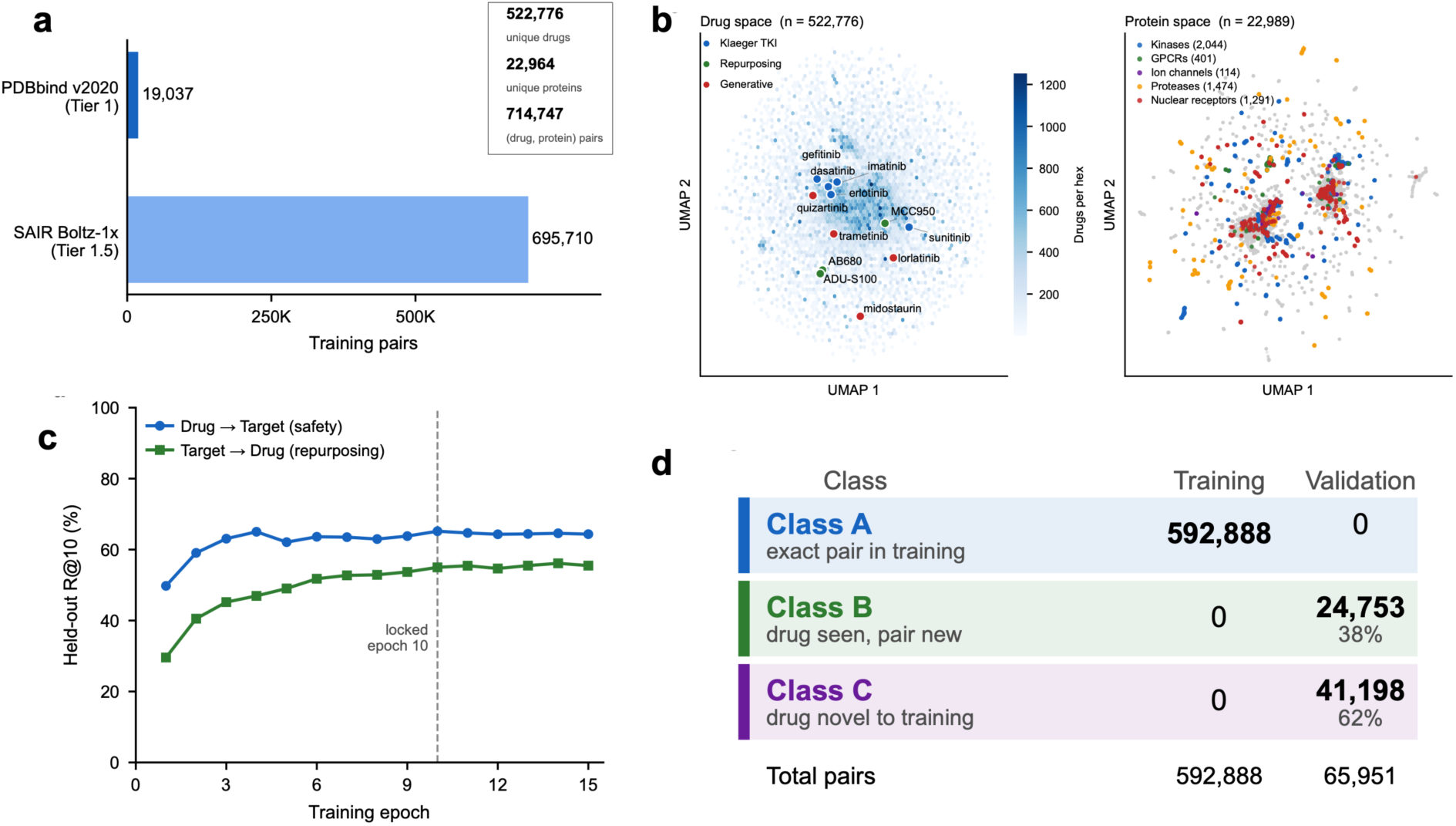
Training corpus, embedding-space coverage, training convergence, and the leakage taxonomy. **(a)** The 714,747 curated (drug, protein) pairs derive from two sources: PDBbind v2020 (19,037 pairs with experimentally measured Kd/IC50)^25^ and SAIR (695,710 pairs from the measured ChEMBL/BindingDB potencies with Boltz-1x predicted interfaces, retained at iPTM ≥ 0.70 and confidence score ≥ 0.70)^26^, spanning 522,776 unique drugs and 22,964 unique proteins; SAIR contributes 37× more pairs than PDBbind. **(b)** UMAP projections of the two embedding spaces. Left: all 522,776 MoLFormer-XL drug embeddings, with anchor compounds overlaid and coloured by section (kinase-inhibitor safety panel, repurposing anchors, generative-design leads); compounds absent from the training library are placed at their nearest ECFP4-Tanimoto training neighbor. Right: all 22,964 ESM-2-650M protein embeddings (mean-pooled over residues), coloured by gene family — kinases, proteases, and nuclear receptors form coherent neighbourhoods. **(c)** Training convergence: held-out validation R@10 over epochs for both retrieval directions; drug→target plateaus at ≈65% by epoch 4–5 and target→drug climbs to ≈55% by epoch 10, where the production checkpoint is locked (dashed line). **(d)** Leakage taxonomy. The split holds out entire MMseqs2 protein-sequence clusters, so every validation/test protein is novel (protein-identity leakage = 0). Each evaluated (drug, protein) pair is graded **Class A** (exact pair seen in training; memorisation), **Class B** (drug seen but never against this protein; drug-level generalisation), or **Class C** (drug novel to training; chemistry novelty). The training split is 100% Class A by construction (592,888 pairs); the validation split contains zero Class A pairs and divides into Class B (24,753; 38%) and Class C (41,198; 62%). The chemical-novelty structure of the Class C drugs is characterized in **Fig. S2** and the direct leakage measurement in **Fig. S1**.

**Table 1.**
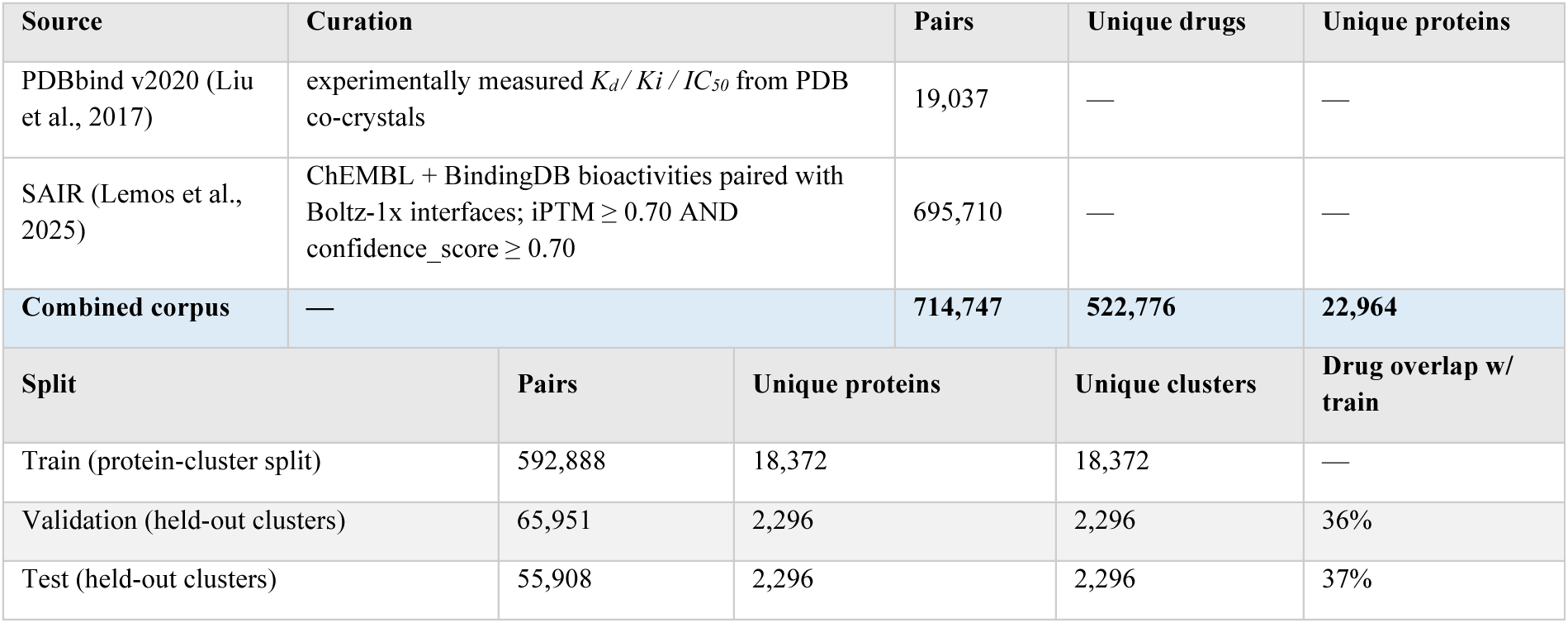
Training data sources and split sizes. The combined dtSFM corpus pairs PDBbind v2020^25^ experimental complexes with SAIR^26^ Boltz-1x-predicted interfaces filtered at iPTM ≥ 0.70 and confidence_score ≥ 0.70. Protein-cluster split by MMseqs2 at 80% sequence identity, with whole clusters assigned to a single partition.

The two pre-trained encoders place the corpus in an organized embedding space (**Fig. 2b**). A UMAP projection of all 522,776 MoLFormer-XL drug embeddings forms a dense, continuous chemical manifold within which the compounds analysed in later sections fall. The 22,964 training proteins separate by gene family: kinases, proteases, and nuclear receptors form coherent neighbourhoods, with G-protein-coupled receptors and ion channels also resolved. The protein encoder therefore organises targets by family before any drug–target training, and this coverage defines what is in-distribution (ID) and out-of-distribution (OOD) for the evaluations below.

Training is monitored by recall-at-K (R@K) on the held-out validation set, which is out-of-distribution in the protein by construction (below). Retrieval is full-pool: for each of the 3,968 validation pairs, the drug→target direction ranks the true target among all 311 unique held-out proteins, and the target→drug direction ranks the true compound among all 3,948 unique held-out drugs, against random-chance R@10 of 3.2% and 0.25%, respectively (**Fig. 2c**). Held-out R@10 rises within the first epochs and plateaus early: the drug→target direction reaches ∼65% by epoch 4–5, while the target→drug direction, which searches the far larger drug pool, climbs to ∼55% by epoch 10; at that checkpoint the median rank of the true partner is 3 of 311 (drug→target) and 6 of 3,948 (target→drug). This early plateau — generalisation reached in the first tenth of training — is the convergence signature expected when an SFM estimates the parameters of a known governing equation rather than fitting an arbitrary function^2,3^. The checkpoint at epoch 10 is locked, early enough to pre-empt memorisation and late enough that both directions have plateaued and is used for every result that follows.

Every evaluation is governed by a two-axis leakage control (**Fig. 2d**), because a retrieval model can inflate its scores by recognising near-duplicates of training examples. The protein axis defines in-versus out-of-distribution: splits assign entire MMseqs2 sequence-identity clusters^29^ to a single partition, so that no validation or test protein — nor any protein closely similar to one — appears in training; direct pairwise measurement confirms zero protein and zero MMseqs2-cluster overlap between training and both held-out splits (**Fig. S1**). The drug axis grades each evaluated (drug, protein) pair by its relationship to training: Class A, the identical pair was seen (memorisation); Class B, the drug was seen but never against this protein target (drug-level generalisation); Class C, the drug is novel to training (chemistry novelty). Because holding out whole protein clusters removes every exact pair, the training split is 100% Class A (592,888 pairs) while the validation split contains no Class A at all, dividing into Class B (24,753 pairs; 38%) and Class C (41,198 pairs; 62%); the full curated set partitions into 592,888 training, 65,951 validation, and 55,908 test pairs. The novelty of Class C is by exact-SMILES identity rather than scaffold: across a 5,000-drug subsample of each held-out split, the maximum ECFP4 Tanimoto similarity of a held-out drug to its nearest training drug has a median of ≈0.69, with ∼37% of held-out drugs near-duplicates of a training compound (Tanimoto 0.9–1.0) and only ∼1–2% having no close training analogue (< 0.3) (**Fig. S2**). Chemistry novelty is therefore reported descriptively, without claiming a hard scaffold-level split; throughout the paper, “ID/OOD” refers to the protein axis and “Class A/B/C” to the drug axis.

### Validating the dtSFM encoder across four output heads, in and out of distribution

We evaluate the locked epoch-10 checkpoint across all four heads in three regimes: ID pairs from protein clusters seen in training, and two OOD sets — a validation and an independent test split — from clusters held out entirely (**Fig. 2d**). Each regime comprises 3,968 pairs, and the performance dashboard consolidates every head across the three regimes (**Fig. 3a**). Retrieval is full-pool and bidirectional: drug→target ranks the true protein among all unique proteins in the regime (928 ID, 311 validation, 349 test), and target→drug ranks the true compound among all unique drugs (≈3,950 throughout), with recall reported as 95% bootstrap confidence intervals over 1,000 query resamples.

**Figure 3.**
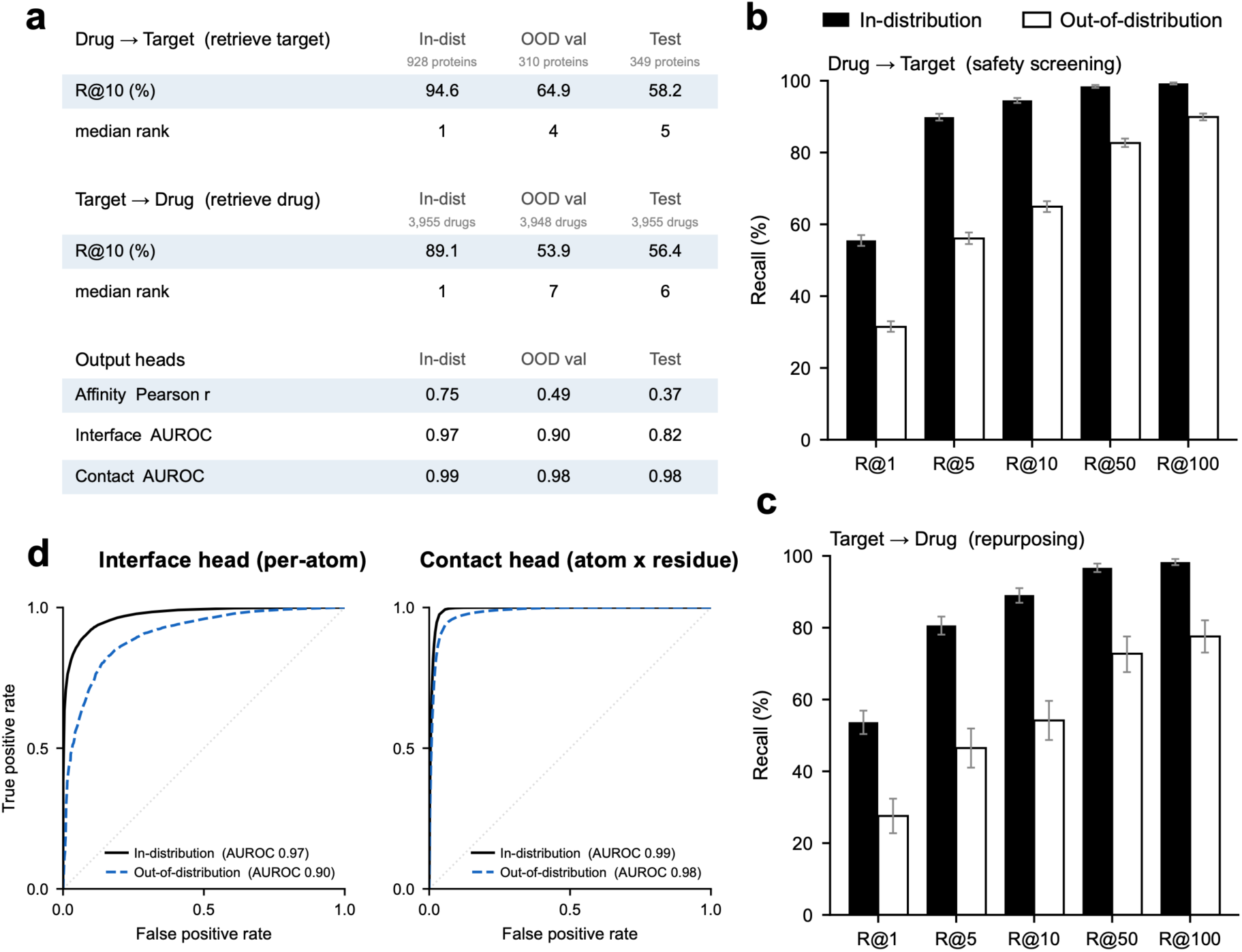
dtSFM encoder validation. In-distribution (ID) evaluation pairs are drawn from protein-sequence clusters seen during training; out-of-distribution (OOD) pairs from clusters held out entirely (MMseqs2 cluster split). Values are the locked epoch-10 checkpoint, ∼4,000 sampled pairs per split. **(a)** Performance dashboard across ID, OOD-validation, and held-out test clusters at epoch 10: retrieval R@10, median rank, and interface/contact AUROC. The affinity head is preliminary and reported in **Fig. S3**. **(b)** Drug→target retrieval (safety-screening direction): recall-at-K (R@1/5/10/50/100) for retrieving a drug’s true target from the unique-protein pool, ID (solid) vs OOD (open); error bars are 95% bootstrap confidence intervals (1,000 resamples). **(c)** Target→drug retrieval (repurposing direction): same convention. **(d)** Interface head (per-atom) and contact head (atom × residue) ROC curves with AUROC; interface AUROC = 0.97 (ID) / 0.90 (OOD), contact AUROC = 0.99 (ID) / 0.98 (OOD) — the contact head generalises almost without loss, while interface and retrieval show the expected ID→OOD gap.

Drug→target retrieval reaches R@10 = 94.6% (ID) and 64.9% (OOD validation), with R@1 = 56% and 32% (**Fig. 3b**). Random retrieval from a pool of N candidates gives R@K = K/N, so random chance is R@1 = 1/928 = 0.1% and R@10 = 1.1% against the in-distribution protein pool (R@10 = 3.2% random chance against the 311-protein OOD pool). Target→drug retrieval reaches R@10 = 89.1% (ID) and 53.9% (OOD), with R@1 = 54% and 28%, respectively (**Fig. 3c**; random R@1 = 0.03%, R@10 = 0.25%). The in-distribution advantage is expected and is larger in the drug→target direction; the ID protein pool (928) exceeds the OOD pool (311), so the in-distribution numbers come from a larger, not smaller, candidate set. For OOD, the median rank of the true partner is 3 of 311 (drug→target) and 6 of 3,948 (target→drug).

The interface and contact heads localise binding on the protein, and the affinity head regresses a scalar binding-strength prediction. The contact head (atom × residue) reaches AUROC 0.99 (ID) and 0.98 (OOD); the interface head (per-atom interface membership) 0.97 (ID) and 0.90 (validation), falling to 0.82 on the test split (**Fig. 3d**). The asymmetry of the contact problem explains why these heads succeed: the ligand side is fully specified — a small molecule is interface in its entirety — so the task reduces to locating the binding residues within the protein sequence, which the PDBbind and SAIR structural labels supervise directly. Contact geometry, being local, transfers across protein families nearly intact, while interface membership, which depends on protein context, is the most sensitive to protein novelty. The affinity head is reported as preliminary (Pearson r = 0.75 ID; 0.48 and 0.37 OOD; **Fig. S3**).

### dtSFM predicts clinical kinase-inhibitor off-targets at proteome scale

The first application was off-target safety screening, in the drug→target direction. dtSFM is operationally simple to use: the user supplies the drug’s SMILES string and a set of candidate target proteins as amino-acid sequences, and dtSFM returns the proteins ranked by predicted binding compatibility. No structural input is required for either partner, and the screening pool can be any user-defined protein set. For the analyses here the pool for each query drug was the 22,964 proteins of the training corpus, which allowed us to apply the leakage taxonomy of **Fig. 2d** directly to every screened pair. dtSFM encoded the drug’s SMILES with MoLFormer-XL, encoded each protein with ESM-2 (the protein encodings are precomputed once and cached), and computed a retrieval cosine for every (drug, protein) pair in the shared embedding space. Proteins mapping to the same gene were collapsed to that gene’s maximum cosine, producing a ranked list of 4,910 genes per drug. Gene rank was the actionable readout: the order in which a laboratory would screen predicted off-targets. We validated this readout against Klaeger et al^5^, a chemoproteomic study in which immobilised broad-spectrum inhibitors capture the expressed kinome from cell lysate, and competition with a free drug, quantified by mass spectrometry, measures that drug’s binding across the captured kinases. The assay is rigorous but bounded: it profiled 243 clinical kinase inhibitors against the roughly 520 kinases the kinobeads matrix can capture, at considerable experimental effort.

dtSFM ranked each drug against all 4,910 genes in its training set, roughly tenfold more targets, and not restricted to kinases. Each (drug, off-target gene) pair was graded by the leakage taxonomy of **Fig. 2d**: Class A (pair in training, i.e. memorisation), Class B (drug and gene each seen but never as this pair), and Class C (drug or gene out-of-distribution). For the nine inhibitors we assembled 57 documented off-target kinases curated from Klaeger and the prior literature. Of these, 34 fell within the kinome Klaeger measured and 23 lay outside it (**Fig. 4a**). Where Klaeger had a measurement, dtSFM rank agreed with it: the documented off-targets the assay scored as binders were those dtSFM ranked near the top (shared block). For the 23 documented off-targets outside the kinobeads matrix — among them ERBB2/4, KDR, FLT1/4, and BLK — the assay was blank, yet dtSFM still ranked them (extension block), recovering known off-targets that the reference experiment could not reach. Under the strict no-leakage Class B condition, dtSFM recovered the documented off-targets at the top of the screen (**Fig. 4b**): across five inhibitors (dasatinib, erlotinib, gefitinib, imatinib, sunitinib), the 27 Class B pairs were recovered at 22.2% within the top 10 of the 4,910-gene screen (6/27), 70.4% within the top 50 (19/27), 88.9% within the top 100 (24/27), and 100% within the top 500; the median Class B off-target ranked 30th of 4,910 genes, the top 0.6% of the screen. dtSFM’s ranking also separated, for each drug, the kinases Klaeger measured as binders from those it measured as non-binders. Scored by AUROC over the 276 kinases common to all nine inhibitors, the separation averaged 0.78 (**Fig. 4c**). It held for compounds whose chemistry never entered training: ibrutinib led all nine inhibitors at 0.94, followed by ponatinib (0.77) and crizotinib (0.76), the three drug-out-of-distribution inhibitors; the in-training inhibitors ranged from 0.61 (imatinib) to 0.84 (dasatinib). A novel compound topping the panel was consistent with the out-of-distribution behaviour seen in encoder retrieval. We tested dtSFM’s top ibrutinib predictions with AF3, which reports interface confidence as the interface predicted TM-score (iPTM, 0 to 1, higher better) and the interface predicted aligned error (PAE) in ångströms (lower better); we treated an interface as confident at iPTM ≥ 0.7 and PAE ≤ 5 Å, and as clinical-grade at PAE ≤ 2 Å. Ibrutinib was itself drug-OOD (Class C). Two of its three highest-ranked cofolds were off-targets documented in the literature but absent from Klaeger’s kinobeads panel: BLK (dtSFM rank 1; iPTM 0.97, PAE 0.82 Å) and JAK3 (dtSFM rank 13; iPTM 0.85, PAE 4.10 Å). The third, ERBB3 (dtSFM rank 9; iPTM 0.97, PAE 1.12 Å), was not a previously annotated ibrutinib off-target, and is a prospective prediction with no wet-lab measurement to confirm or refute it. BLK and ERBB3 reached clinical-grade interfaces; JAK3 was confident but weaker.

**Figure 4.**
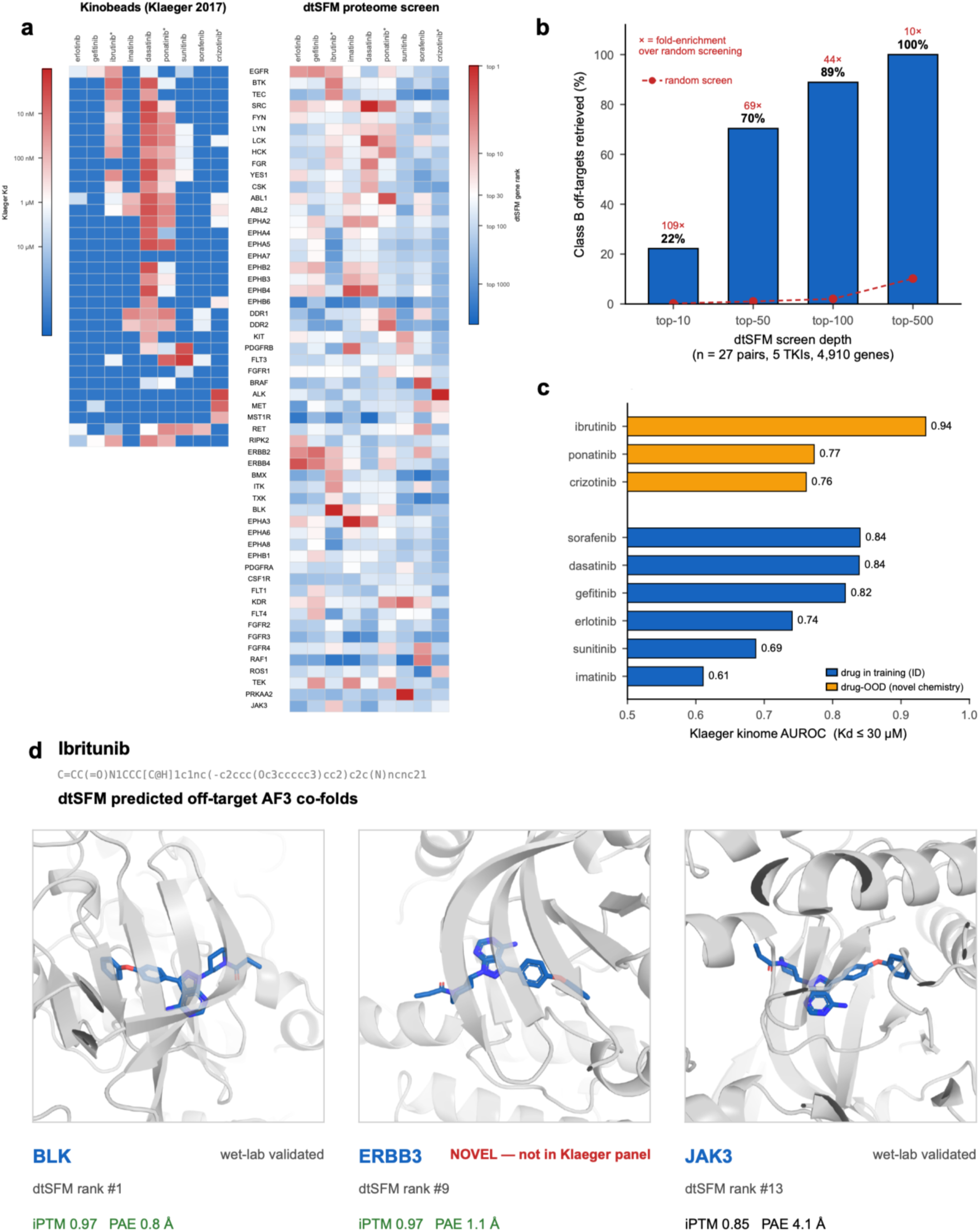
Proteome-scale off-target safety screening, validated against a published chemoproteomic kinase panel. In the drug→target direction, dtSFM ranks all ≈4,910 unique training genes for each drug by best drug–protein cosine; gene rank is the actionable readout. Each (drug, off-target gene) pair is graded by the leakage taxonomy of Fig. 2d. **(a)** Concordance heatmap: Klaeger et al.^5^ kinobeads measured affinity (left) vs dtSFM gene rank (right) for nine clinical kinase inhibitors × named off-targets grouped by kinase family; the shared block (the kinome Klaeger quantified) and the extension block (documented off-targets outside the kinobeads matrix — ERBB2/4, KDR, FLT1/4, BLK) show dtSFM recovers the measured pattern and continues proteome-wide. **(b)** Cumulative recovery of the 27 Class B (drug, off-target) pairs across five inhibitors at top-K of the 4,910-gene screen — 22.2% (top-10), 70.4% (top-50), 88.9% (top-100), 100% (top-500); median Class B rank 30 of 4,910 (top 0.6%). **(c)** Per-drug zero-shot AUROC for separating each inhibitor’s Klaeger binders from non-binders over the 276-kinase comparison set; mean AUROC 0.78. **(d)** AF3 structural verification of ibrutinib’s top dtSFM off-target predictions.

### dtSFM virtually screens a half-million-compound library against immunology targets

The second application was virtual screening for drug discovery and repurposing, in the target→drug direction. For each target, dtSFM computed the retrieval cosine: the thermodynamic binding-compatibility score the SFM produces from sequence, between the target and every one of the 522,776 unique compounds in the v3 training library, returning a fully ranked library in a single pass. The three targets were NLRP3, CD73, and STING1, immunology proteins with active drug-discovery programs in inflammatory disease and cancer immunotherapy ^30–32^. They were also chosen to span the leakage-class spectrum, which let us probe model behaviour across training-coverage regimes within a single application: CD73 is heavily represented in training (Class A-dominant top hits), STING1 is sparsely represented (Class B-dominant top hits), and NLRP3 sits between them.

The library screen separated a small high-cosine tier from a broad low-cosine bulk. Across the three targets the median library cosine sat near zero (NLRP3 0.04, CD73 −0.01, STING1 0.03), while only the top 1,000 compounds of the 522,776 reached cosines of 0.21–0.29, the top 100 reached 0.31–0.35, and the per-target maximum was 0.35–0.41. Published anchor compounds ranked within this top tier in proportion to their leakage class (**Table 2**). Three of the Class B anchors below (MCC950 for NLRP3, AB680 and AOPCP for CD73) were trained against the mouse ortholog of their human target only, so their Class B assignment is cross-species rather than same-species pair-novelty. For NLRP3, the Class B canonical inhibitor MCC950 ^33^ ranked 3 of 522,776 (cos 99.999%); the Class A approved drug Glyburide (repurposed for NLRP3 activity) ranked 2,936 (cos 99.4%); and the Class B approved drug Tranilast ranked 6,635 (cos 98.7%). For CD73, the Class B gold-standard inhibitor AB680 ^34^ ranked 503 (cos 99.9%), the Class A clinical-stage LY3475070 ranked 1,118 (cos 99.8%), and the Class B substrate analogue AOPCP ranked 1,276 (cos 99.8%). STING1 had no clinical anchors in the v3 training library, all known agonists being Class C; when their dtSFM cosines against the STING1 target embedding were computed and inserted into the 522,776-compound distribution, ADU-S100 ^35^ reached rank 913 (cos 99.8%), MK-1454 rank 5,698 (cos 98.9%), and SR-717 rank 3,956 (cos 99.2%).

**Table 2.**
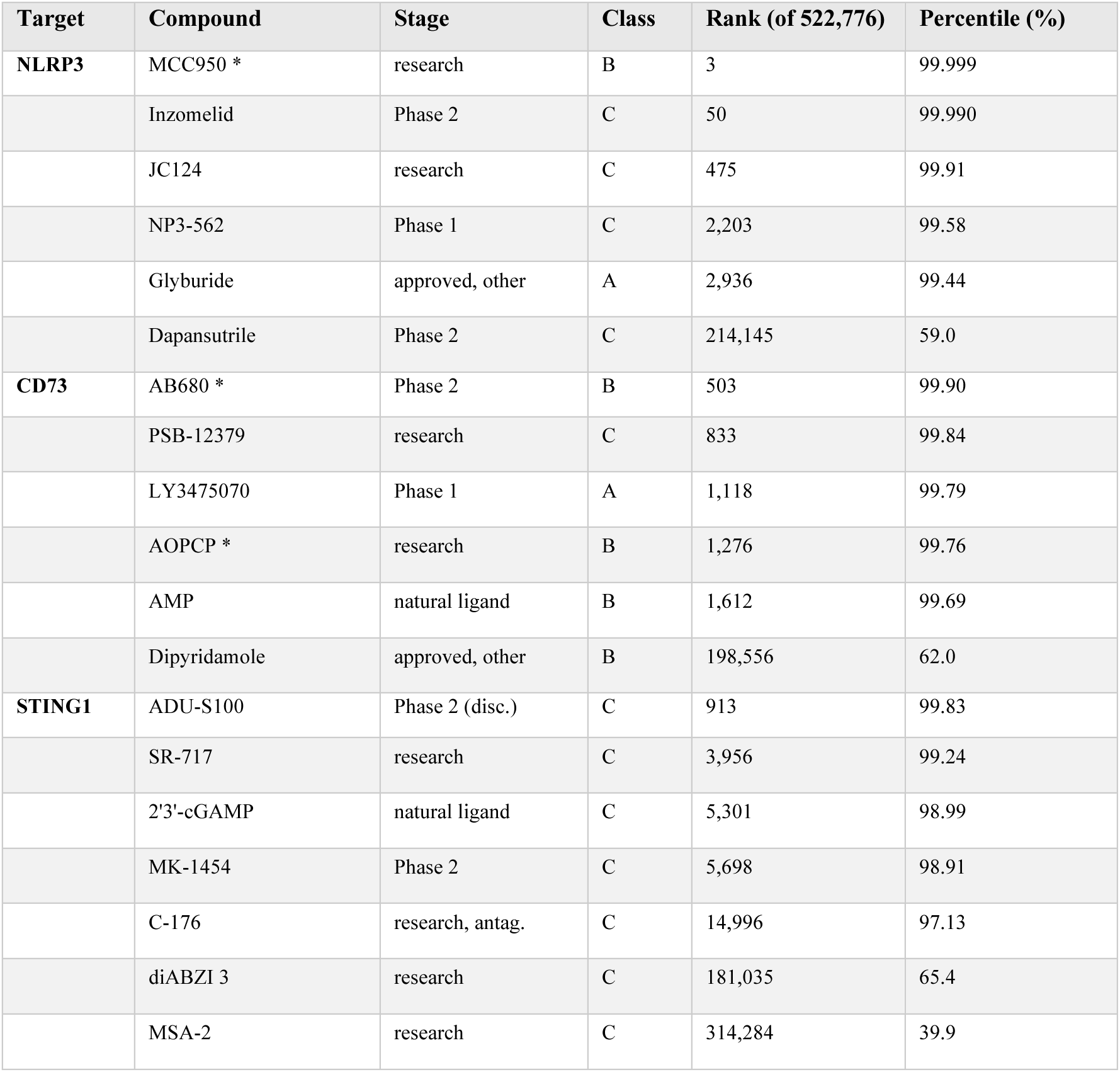
Published anchor compounds recovered by dtSFM cosine ranking against the 522,776-compound library for NLRP3, CD73, and STING1. All three targets are represented in training, so leakage class is defined by the drug’s relationship to the training set: A = the (drug, target) pair was seen in training (memorisation upper bound); B = the drug was seen in training but never paired with this target (genuine repurposing prediction); C = the drug was never seen in training (drug-out-of-distribution; novel chemistry). *Target represented in training only via its mouse ortholog, so the human-target pairing is cross-species — these compounds are Class B because the drug was seen against the mouse ortholog, not the human target.

For AF3 verification of the dtSFM screen, we drew per-target samples spanning the dtSFM-prioritized top of each ranking together with matched controls. From each target’s library ranking we took the top-50 dtSFM-cosine hits, a scaffold-diverse sample of 35–50 next-tier hits, the published anchor compounds for that target (5–10 each, Class A/B/C), and 5 matched negative controls. The per-target totals were 95 pairs for CD73, 115 for NLRP3, and 113 for STING1, summing to 323 pairs, and AF3 was run as a cofold on every pair. Plotted as dtSFM cosine percentile against AF3 iPTM (**Fig. 5a**), most compounds in the dtSFM-prioritized strata cleared the AF3 binder floor of iPTM 0.7. In the top 0.01% of the library by dtSFM cosine (rank ≤ 52, 156 pairs across the three targets), the bulk sat above the AF3 iPTM 0.7 line, including most published anchors. Three Class C clinical compounds illustrated complementarity between the two models. Dapansutrile (NLRP3 phase-2 inhibitor), MSA-2 (STING1 agonist), and diABZI 3 (STING1 agonist) are absent from the v3 training library; when dtSFM computed their cosines against their respective targets and inserted them into the 522,776-compound distribution, Dapansutrile reached cos 59% (NLRP3), MSA-2 cos 40% (STING1), and diABZI 3 cos 65% (STING1), all well below the top-cosine tier. AF3 nonetheless placed them at confident interfaces. dtSFM did not flag these compounds as top-cosine candidates for their targets, while AF3 confirmed binding.

**Figure 5.**
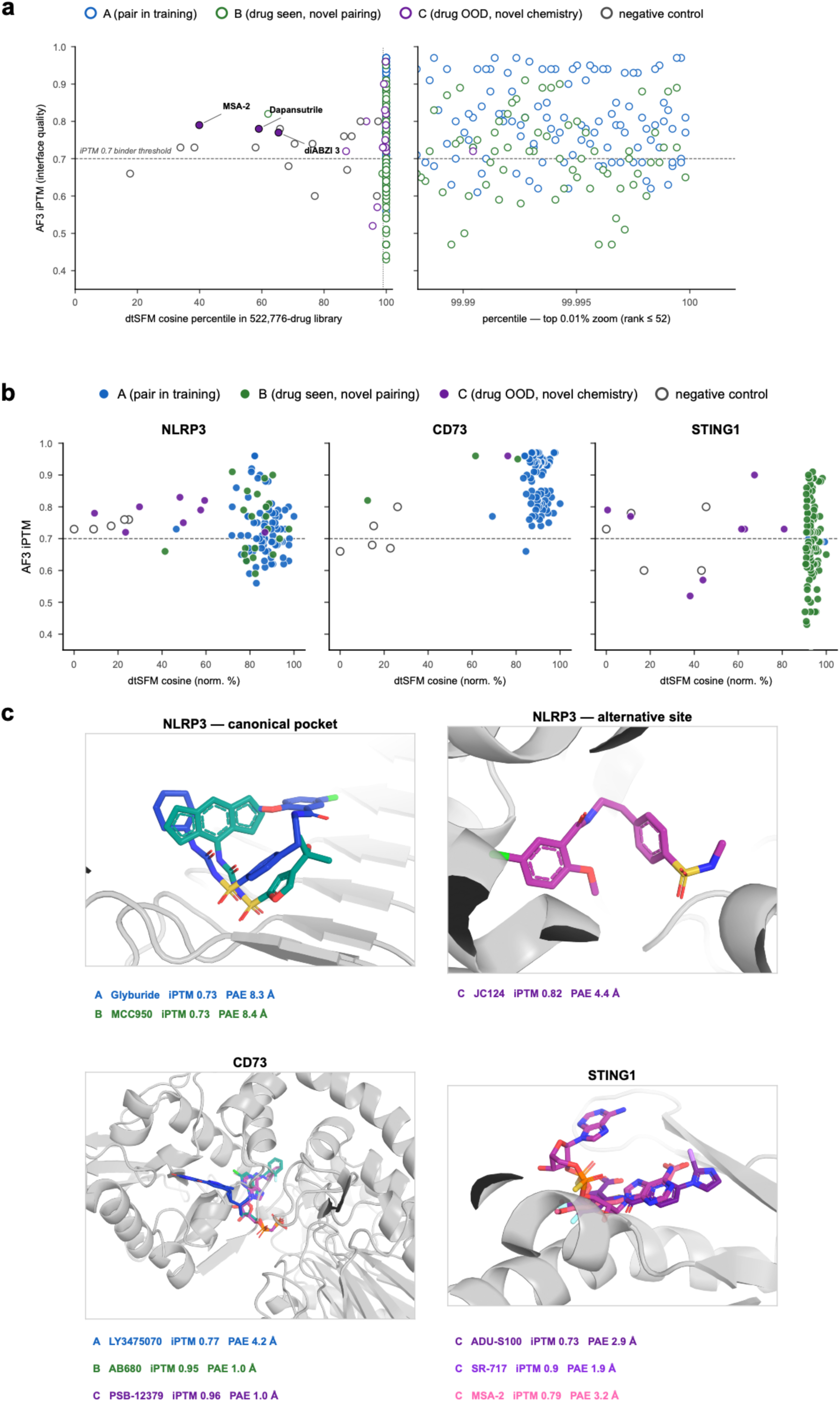
Virtual screening and repurposing across three immunology targets. For each target, dtSFM computes the retrieval cosine between the target sequence and every one of the 522,776 unique training-library compounds, returning a fully ranked library in a single pass; NLRP3, CD73, and STING1 were chosen to span the leakage-class spectrum (CD73 Class A-dominant, STING1 Class B-dominant, NLRP3 intermediate). A 323-pair verification cohort (top-50 cosine hits, scaffold-diverse next-tier hits, published anchors, and matched negative controls) was cofolded by AF3. **(a)** Screen × structure complementarity: dtSFM cosine percentile vs AF3 iPTM (full library and top-0.01% zoom), coloured by leakage class; three Class C clinical compounds (dapansutrile, MSA-2, diABZI compound 3) fall below the top-cosine tier yet AF3 places each at a confident interface. **(b)** Per-target class composition: CD73 top hits are training-saturated (Class A), STING1 top hits are genuine novel predictions (Class B) clustering near the binder floor, and NLRP3 is mixed. **(c)** AF3-based structural characterization at the canonical druggable pockets, with iPTM and interface PAE annotated per compound.

Per-target class composition (**Fig. 5b**) made this regime-dependence explicit. Each panel plots the target’s full AF3 verification cohort as dtSFM cosine percentile against AF3 iPTM. The leakage spectrum was sharpest at the top of each ranking. CD73’s top-50 cosine hits were entirely Class A (50/50): every highest-cosine prediction was a (compound, target) pair the model had already seen, giving clean validation but no novel repurposing leads. STING1 was the inverse — 48 of its top-50 hits were Class B (genuine novel predictions, the drug seen in training but against a different target); the AF3 confidences of these STING1 hits ranged from above and below binder floor (mean top-50 iPTM 0.70) rather than well above it, so the predictions were novel but only moderately structurally supported. NLRP3 sat between the two: its top-50 was training-heavy (42 Class A / 8 Class B). NLRP3 also produced the cleanest published-anchor result: all 8 Class C anchor compounds and 1 of 2 Class B anchor compounds cleared the AF3 binder gate. Across the top-50-cosine and scaffold-diverse strata of the three targets, 46 novel Class B/C compounds cleared the combined AF3 binder gate (iPTM ≥ 0.7 AND interface PAE ≤ 5 Å): 38 for STING1, 8 for NLRP3, and 0 for CD73, the last consistent with its top being saturated by training pairs.

AF3 placed dtSFM’s hits at the canonical druggable pockets of each target (**Fig. 5c**), with interface confidence that varied by target. CD73 cofolds were the most confident, AB680 (iPTM 0.95, PAE 0.98 Å) and PSB-12379 (iPTM 0.96, PAE 1.02 Å) at near-clinical-grade interfaces and LY3475070 within the binder gate (iPTM 0.77, PAE 4.16 Å). STING1 cofolds also cleared the binder gate, SR-717 the strongest (iPTM 0.90, PAE 1.89 Å), followed by MSA-2 (iPTM 0.79, PAE 3.16 Å) and ADU-S100 (iPTM 0.73, PAE 2.88 Å). NLRP3 was the weaker case: Glyburide (iPTM 0.73, PAE 8.31 Å) and MCC950 (iPTM 0.73, PAE 8.38 Å) were placed in the canonical NACHT pyrin pocket but at low interface confidence, failing the combined binder gate on interface PAE despite being known NLRP3 binders, while JC124 sat in an alternative sub-pocket and cleared the binder gate (iPTM 0.82, PAE 4.42 Å). The pipeline thus surfaced both the canonical and an alternative NLRP3 druggable surface, with the alternative pocket better structurally supported than the canonical one for these compounds.

### dtSFM generates novel drug candidates that AlphaFold-3 confirms at scale

The third application was generative design, in the target→drug direction extended beyond any existing library. A decoder is a generative model that produces new sequences token by token (a token is a short symbol from the SMILES alphabet, typically a single character such as’C’,’N’, or’=’). dtSFM’s decoder is a cross-attentive autoregressive Transformer that generates the SMILES string of a novel compound one token at a time while attending to the per-residue features of a target protein. It follows the autoregressive language-model paradigm developed for biological-sequence generation, which has been applied to protein sequences in ProGen^36^ and to SMILES in MolGPT^37^; what is new here is the cross-attention conditioning on the dtSFM encoder’s target embedding, which yields target-specific molecular generation from the same physics-derived embedding space used for retrieval. It is the first trained instance of the cross-attentive SFM decoder specified for antibody-antigen^21^ and other molecular recognition systems^2^. The dtSFM decoder was trained on the same 714,747 (drug, target) pairs as the encoder by teacher-forced cross-entropy on the ground-truth SMILES, with an auxiliary loss that pulls each generated molecule’s embedding into the encoder’s preferred basin for its target. Base sampling performance (RDKit validity, reconstruction Tanimoto to the ground-truth drug, encoder cosine to the conditioning target, QED and Lipinski-Ro5 compliance) is reported in **Fig. S4**. We applied the decoder to 16 immunology and oncology targets, generated 1,200 candidate molecules (decoder checkpoint *decoder_v0.2_step50K* over encoder epoch_010), and verified every candidate by AF3 cofolding. The full generative cascade has five stages (**Fig. 6a**): target-conditioned decoder generation; dtSFM-encoder reranking against the intended target with a max-Tanimoto-to-approved ≤ 0.40 chemistry-novelty filter; AF3 cofolding for on-target verification; off-target safety screening across the human proteome; and AF3 paralog selectivity cofolding.

**Figure 6.**
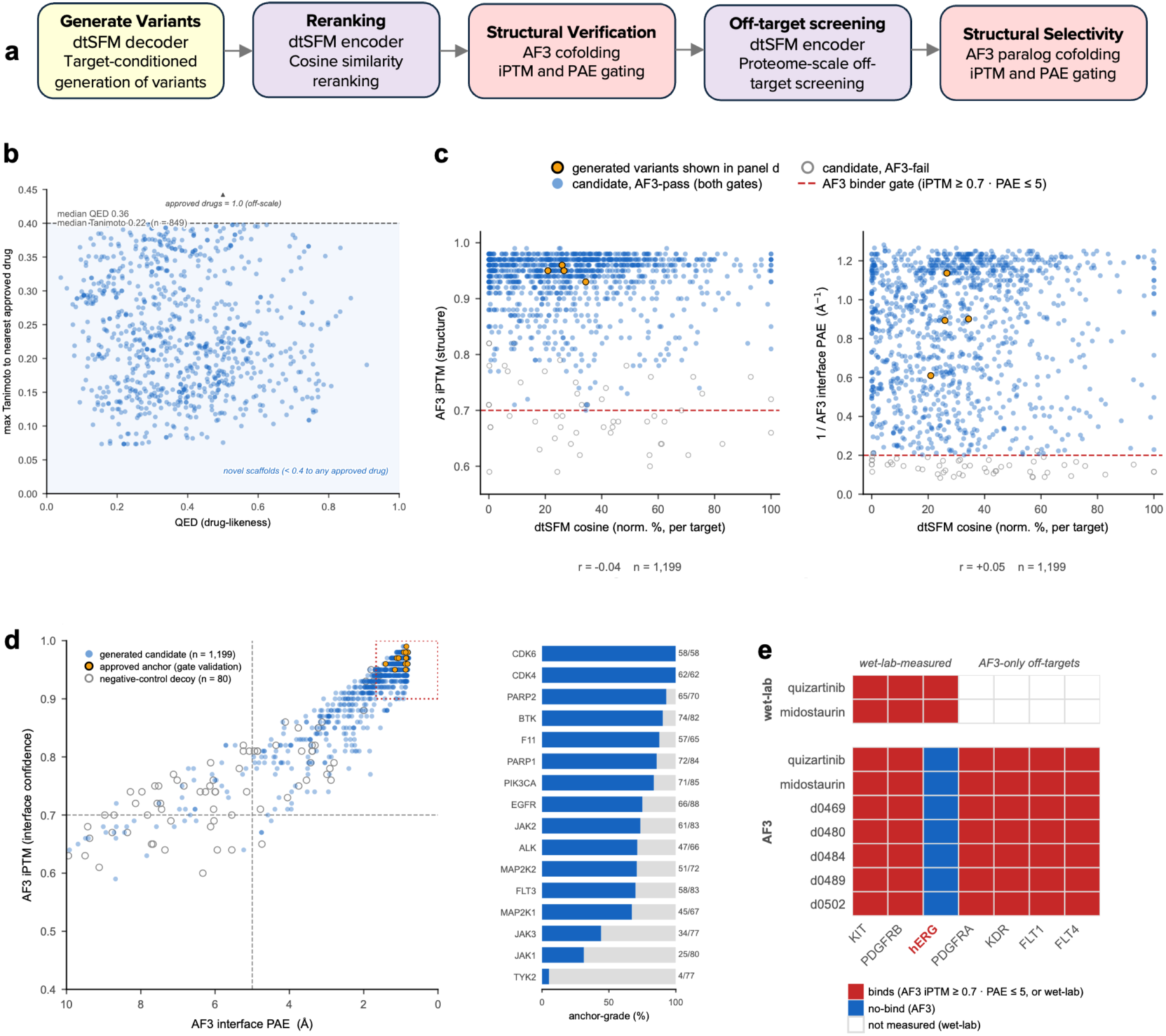
The dtSFM decoder generates novel candidates that AlphaFold 3 confirms across 16 targets. The first trained instance of the cross-attentive SFM decoder, applied to 16 immunology and oncology targets, generated 1,200 candidate molecules, each verified individually by AF3 cofolding. **(a)** The five-stage generative cascade: target-conditioned decoder generation; dtSFM-encoder reranking with a max-Tanimoto-to-approved ≤ 0.40 chemistry-novelty filter; AF3 on-target cofolding; proteome-wide off-target safety screening; and AF3 paralog selectivity cofolding. **(b)** Generation quality (n = 849 reranked candidates): ECFP4 Tanimoto to the target’s nearest approved drug vs the QED; every reranked candidate is novel chemistry relative to the approved drug (max Tanimoto ≤ 0.40, median 0.22) and none duplicates a training compound (median Tanimoto 0.73 to the nearest training compound). **(c)** dtSFM and AF3 are uncorrelated: per-target-normalised dtSFM cosine vs AF3 iPTM (Pearson r = −0.04) and vs 1/interface PAE (r = +0.05). **(d)** AF3 iPTM vs interface PAE (axis reversed) for every generated candidate, the 15 approved-drug anchors, and 80 negative-control decoys; 850 of 1,200 candidates (70.8%) clear the anchor-grade gate (iPTM ≥ 0.9 AND PAE ≤ 1.67 Å), matching the AF3 confidence of the approved drugs, while only 2 of 80 decoys do; per-target anchor-grade fractions at right. **(e)** Binary AF3 binds/no-bind matrix (binds = iPTM ≥ 0.7 AND interface PAE ≤ 5 Å) for the FLT3 series, two approved inhibitors and representative dtSFM designs, against six kinases and the cardiac hERG channel (KCNH2), with measured wet-lab calls shown for KIT, PDGFRB and hERG.

Generation quality is summarised by chemistry similarity by ECFP4 Tanimoto (as in **Fig. 2b** and **Fig. S2**) and drug-likeness by QED, a 0–1 composite of molecular weight, lipophilicity, polar surface area, and related properties; a coarse pre-screen, not a full ADMET profile. Of the 1,200 generated candidates, 849 were retained after the reranking + chemistry-novelty filter (max Tanimoto-to-approved ≤ 0.40). The chemistry-novelty filter excluded 110 of the 1,200 unfiltered candidates (9.2%), whose maximum ECFP4 Tanimoto to the cohort’s approved drug exceeded 0.40, including 8 candidates (0.7%) that were exact matches to training set molecules; the remaining 1,090 were then narrowed to 849 by the encoder rerank and a basic Lipinski-Ro5 / QED drug-likeness screen (**Fig. 6b**). By construction, every one of the 849 reranked candidates therefore had max Tanimoto to its approved drug ≤ 0.40 (median 0.22 across the 849); the median Tanimoto to the nearest training compound was 0.73, indicating that most candidates are novel variants of training chemistry, with a tail of 94 (11%) reaching scaffold-novel (Tanimoto < 0.5) (**Fig. 6b**; **Fig. S4**). The cohort’s QED median was 0.36, modest for a base decoder not optimised for ADMET. A structural-alert audit of the generated cohort found that 568 of 1,200 candidates (47%) carried at least one medicinal-chemistry liability — a reactive sp3 alkyl halide (226), a bare phosphine (218, a non-drug-like decoder artefact), or a free thiol (124) — and many cleared the anchor-grade gate regardless, because AF3 scores pocket complementarity and is blind to whether a terminal group is reactive or synthesizable. Of the 850 anchor-grade designs, 471 (55%) were also free of these alerts.

Before reporting structural confidence, we first establish that dtSFM and AF3 carry independent information about binding, so that any agreement between them is genuine corroboration rather than a restatement of the same signal. The two models differ in every aspect of their construction. dtSFM is sequence-native: amino-acid strings and SMILES as inputs, paired binding annotations as training data, a contrastive cosine loss in a 512-dimensional embedding space, and a frozen-encoder Transformer with cross-attention fusion as the architecture. No three-dimensional coordinates appear anywhere in dtSFM’s training, loss, or inference path; the decoder generates candidates by thermodynamic selection in the embedding space, not by sampling 3D structures. AF3, the verifier we use here, operates in 3D coordinate space and shares no architectural component, training data, loss, or representational primitive with dtSFM. The empirical signature of that orthogonality is uncorrelated scoring (**Fig. 6c**): across the 1,199 generated candidates carrying both metrics, the per-target-normalised dtSFM cosine showed essentially zero Pearson correlation with AlphaFold-3 iPTM (r = −0.04) and with inverse interface PAE (r = +0.05). AlphaFold-3’s confirmation is therefore an independent source of binding information rather than a restatement of the dtSFM cosine signal. This contrasts with the structure-first protein-binder design family (RFdiffusion^13^, BindCraft^14^, BoltzGen^15^), where the generator and verifier share a three-dimensional structural representation.

For structural verification we use two AF3 confidence gates throughout this section, both built on the iPTM and interface PAE metrics. The binder gate (iPTM ≥ 0.7 AND interface PAE ≤ 5 Å) is the broad criterion for a confident interface, the same gate applied in **Fig. 5**. The anchor-grade gate (iPTM ≥ 0.9 AND interface PAE ≤ 1.67 Å) is the stringent criterion; its PAE bound equals the worst PAE among the 15 approved-drug anchors cofolded in this study (so all 15 clear it by construction), meaning a generated candidate clearing the same gate matches the AF3 structural confidence of the approved drugs themselves. Of the 1,200 generated candidates, 1,146 (95.5%) cleared the binder gate and 850 (70.8%) cleared the anchor-grade gate (**Fig. 6d**). For comparison, 28 of 80 negative-control decoy drugs (35%) cleared the binder gate but only 2 of 80 (2.5%) cleared the anchor-grade gate (**Fig. S5**): the binder gate sits at the AF3 drug-like floor that necessitates orthogonal verification, while the anchor-grade gate effectively excludes decoys.

Per-target performance was heterogeneous (**Fig. 6d**, right; **Table 3**). Anchor-grade fractions ranged from 100% (CDK4 62/62, CDK6 58/58) to 5% (TYK2 4/77), with the majority of targets above 70%. The mean AF3 iPTM per target was between 0.79 (TYK2) and 0.96 (PARP2), and 14 of the 16 targets had a mean iPTM ≥ 0.90 with mean interface PAE below 2 Å; the best generated design per target reached iPTM 0.95–0.99 and interface PAE 0.79–1.37 Å, near-crystallographic. This anchor-grade population is shown structurally in **Fig. 7**, where 5 highly ranked designs per target ranked by AF3 iPTM and interface PAE are cealign-overlaid on the approved drug at each target’s binding pocket; the SMILES and computed properties of every shown design are in **Table S1**, and individually in **Fig. S6**. Although these ensembles converge structurally on the approved drug’s pose, the designs are by construction chemically distinct from the approved drug (max ECFP4 Tanimoto ≤ 0.40 by the novelty filter; **Fig. 6b** and **Table S1**). Structural pose convergence (binding the same pocket) and chemical similarity (substructure overlap by Tanimoto) are distinct measures of drug similarity. The decoder produces molecules that are chemically novel relative to the approved drug yet adopt the same binding mode according to AF3.

**Figure 7.**
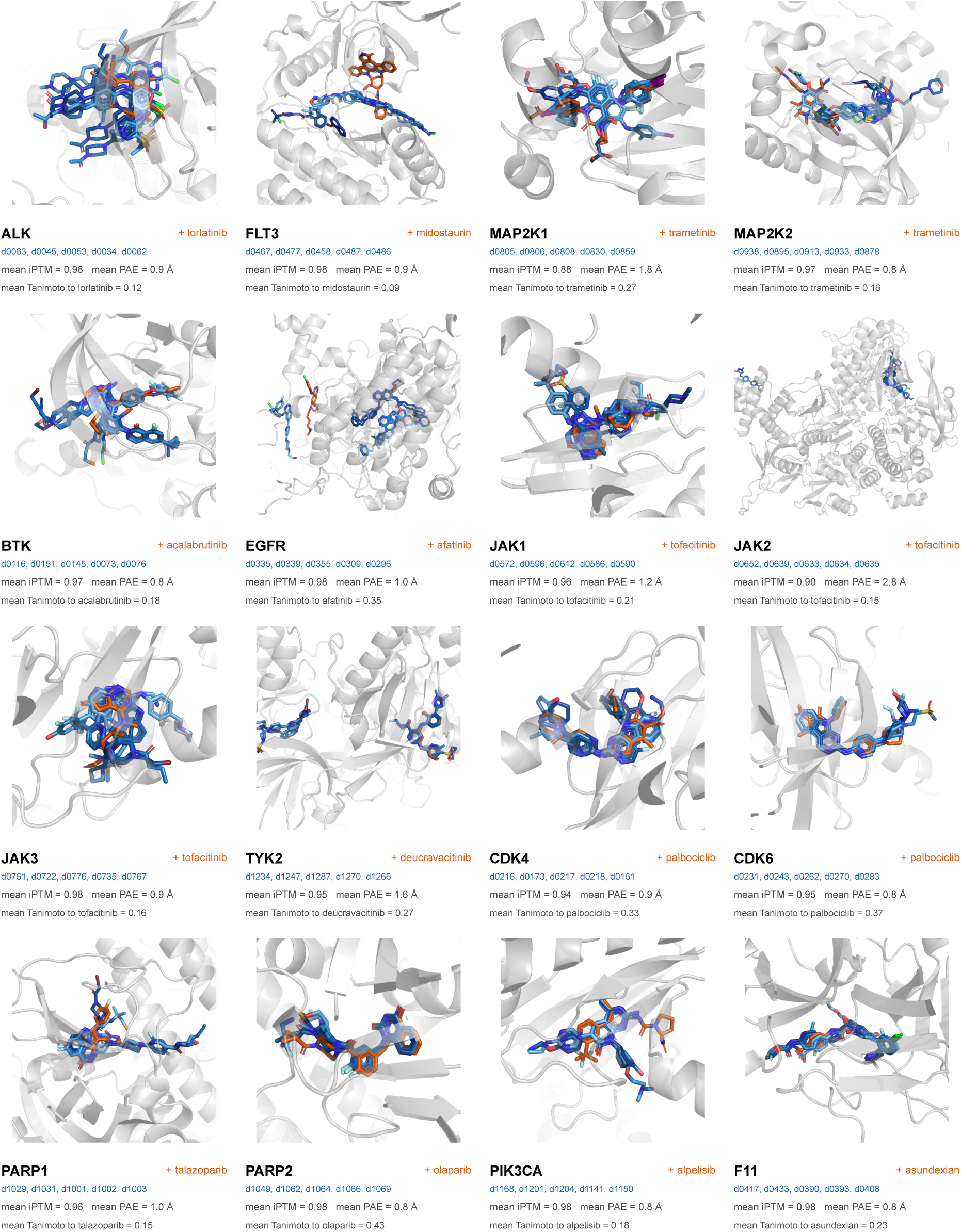
Structural-validation of dtSFM generated designs converge on the approved-drug pocket across all 16 targets. For each target the five of the most AF3-confident generated designs (blue) are overlaid on the approved drug (orange) cofolded against the same target; the protein is shown once as a grey cartoon. Each cell reports the design identifiers (dNNN), the ensemble-mean AF3 confidence (mean iPTM and mean interface PAE), and the mean ECFP4 Tanimoto of the five designs to the approved drug. All structures are AF3 cofolds; SMILES and computed properties of every shown design are listed in **Table S1** and shown individually in **Fig. S6**.

**Table 3.**
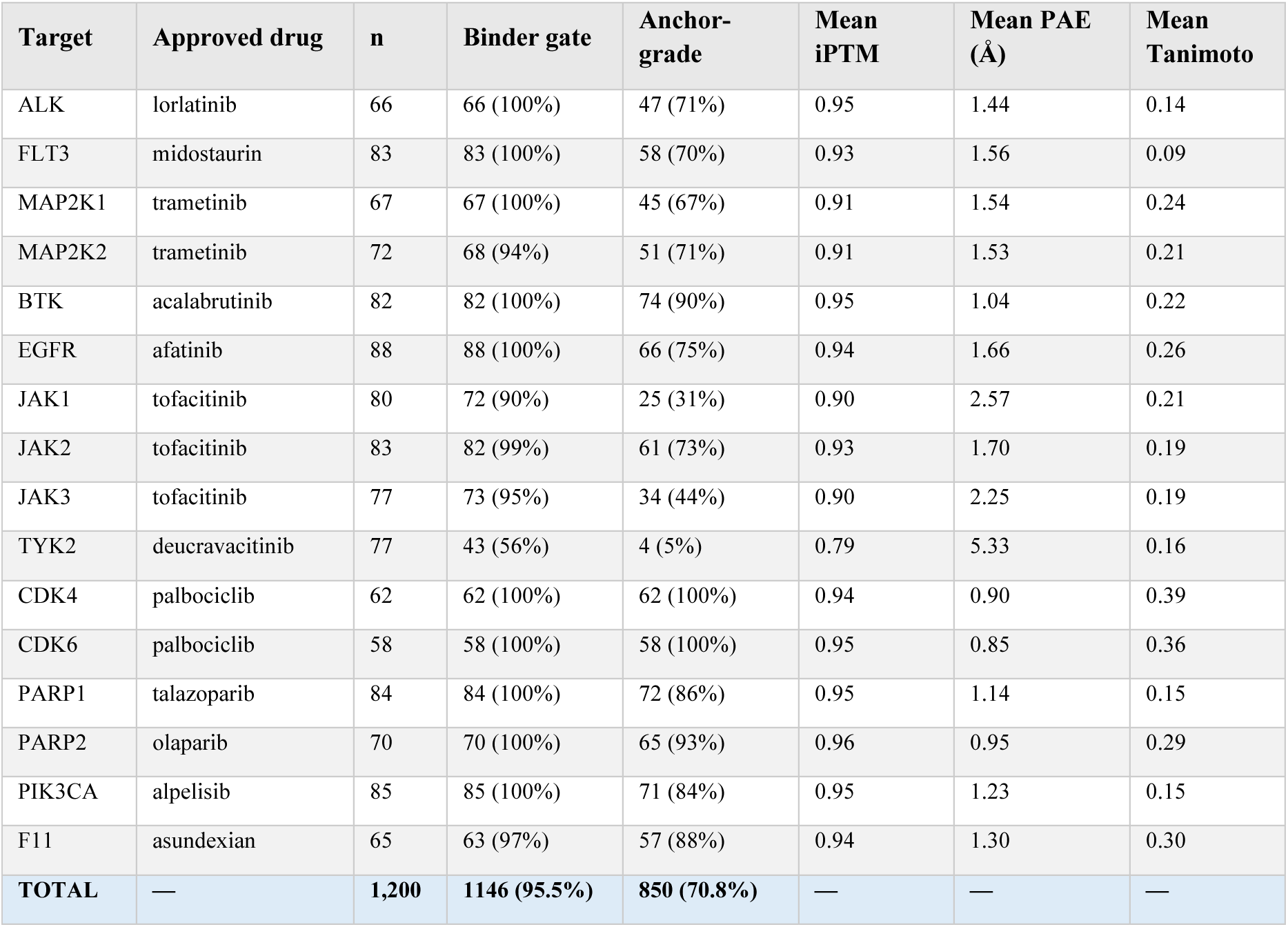
Per-target decoder generation across 16 immunology/oncology targets. Binder gate: AF3 iPTM ≥ 0.7 AND interface PAE ≤ 5 Å. Anchor-grade gate: iPTM ≥ 0.9 AND interface PAE ≤ 1.67 Å (matches the worst-PAE among the 15 approved-drug anchors cofolded in this study). Mean ECFP4 Tanimoto (radius 2, 2,048 bits) computed against the per-target approved-drug anchor across all generated candidates with parseable SMILES.

One limit of this verification arm is AF3’s hERG blind spot (**Fig. 6e**). For the FLT3 series, we cofolded the approved kinase inhibitors quizartinib and midostaurin against their measured kinase targets^5^ and against the hERG potassium channel (KCNH2). AF3 reproduced the wet-lab kinase binding (both drugs at KIT and PDGFRB) but scored hERG as no-bind for every compound tested, including the two approved drugs that clinically block hERG (quizartinib carries an FDA boxed warning for QT prolongation). The dtSFM-generated FLT3 designs fell in the same AF3 no-bind zone for hERG. Any hERG claim for these designs will therefore require wet-lab testing, not AF3-verification.

## DISCUSSION

The dtSFM applies the SFM architecture to small molecule drug–target protein binding, executing off-target safety screening, library repurposing, and de novo generative design from sequence alone. Because the architecture is prescribed by the identity between the transformer softmax attention and the Boltzmann distribution governing molecular binding^1,2^, the model is not approximating an unknown function but estimating the parameters of a known one. Its retrieval cosines are estimates of the partition-function-normalised binding compatibility between drug and target, and the same scores govern both directions of retrieval and the decoder’s sampling distribution. This is computational thermodynamics in practice: the model computes a thermodynamic quantity from sequence, and the same computation supports retrieval, screening, and conditional generation across the public-data scale of drug–target binding, consistent with the Molecular Recognition Computing (MRC) framework and its empirical performance in other binding domains^3,21^. Two properties of drug–target recognition explain why the dtSFM domain scales effectively to a generative decoder. The first is interface density: a small-molecule drug is the maximally interface-dense agent — essentially the entire molecule is the binding interface, in contrast to an antibody (only the CDR loops contact) or an antigen (only the epitope) — which anchors the contrastive signal without the residue-level masking that diffuse partners require. The second is data availability: drug–target binding has been measured for decades, and combining experimental structures with experimental binding data and structural modeling yields 714,747 paired interactions across half a million compounds, sufficient volume to train not only an encoder but a generative decoder.

The orthogonal-verifier construct is the architectural commitment that distinguishes dtSFM from the structure-first generative design family. In the dominant structure-design and structure-verify cycle, structure-first tools generate or optimise novel binder structures in three-dimensional coordinate space: RFdiffusion^13^, BindCraft^14^, and BoltzGen^15^ for protein binders; DiffSBDD^16^, TargetDiff^17^, and Pocket2Mol^18^ for small-molecule ligands. The designs are then validated by structural prediction tools such AlphaFold-class or Boltz-class models. Two couplings make this cycle difficult. First, the generator and the verifier share a three-dimensional representation, so the same structural prior is exposed twice — once in generation and once in validation — which limits how independent the verification can be. Second, the verifier does not score 3D coordinates directly; it requires the structure to be translated back into a discrete representation (a protein sequence for protein binders, a chemically valid molecule for ligands), and the translation step introduces its own sources of error. For protein designs the translation is inverse folding (typically ProteinMPNN^38^), whose outputs can sit outside natural protein sequence distributions and inflate iPTM artefactually without corresponding to physical binding^39^. For small-molecule designs the translation is post-hoc bond inference from generated atom positions; recent benchmarking finds that a small minority of molecules generated across the leading small-molecule generators have structurally valid conformations, with frequent three-membered rings and valence violations^40^. Reported wet-lab hit rates in both regimes commonly remain in the single-to low double-digit percent range, and high AF3 confidence does not consistently predict experimental binding^39^. The dtSFM–AF3 pairing decouples these two arms. dtSFM and AF3 share no architecture, training data, loss, or underlying representation (**Fig. 6c**, r ≈ 0), and the dtSFM decoder emits a SMILES string directly with valid chemistry enforced by the tokenisation grammar; no post-hoc inverse-folding or bond-inference step intervenes. The novel SMILES sample within the distribution of chemistry the encoder and decoder were trained on (median Tanimoto 0.73 to the nearest training compound, only 11% scaffold-novel at Tanimoto < 0.5), and the target protein is a natural sequence drawn from the same public databases AF3 was trained on; both inputs to AF3 therefore sit inside its training distributions. The actor–verifier decomposition specified by the MRC framework^2^ formalises this design principle; dtSFM and AF3 instantiate it for drug–target, and the same pairing of an SFM-based generator with a structure-prediction verifier extends naturally to other binding domains.

The dtSFM-specific failures trace to identifiable properties of the training data, and the model is not static: its supervision is paired binding data augmented with structural labels from co-folding predictors, and both inputs can grow substantially with the next curation iteration. Per-target training density does not predict generative performance monotonically. CDK4 and CDK6 reach 100% anchor-grade from only 2 and 9 training protein entries respectively, while TYK2 falls to 5% from 20 entries, and the four Janus kinase family members (TYK2 5%, JAK1 31%, JAK3 44%, JAK2 73%) underperform jointly despite reasonable training counts (20–68 entries each), indicating a family-level limitation that more entries of the same structural-prediction quality will not fix. A second pattern shows the decoder faithfully sampling the chemical distribution of its training data: F11 (coagulation factor XI) generates candidates that reach 88% anchor-grade but 0 of 65 satisfy the Lipinski Ro5, because F11’s training chemistry is dominated by large, non-Ro5 inhibitors. dtSFM’s path forward follows directly: per-family curation of higher-confidence residue-level structural labels via co-folding (scaling the SAIR strategy), harmonised single-assay supervision for the affinity head, and explicit objectives or constrained sampling when the design goal departs from the training chemistry. dtSFM training supervision can therefore expand continually as the public bioactivity corpus grows and as co-folding generates residue-level structural labels for more pairs, without an architectural change.

Conventional deep-learning models scale their parameter counts with the complexity of the function they approximate. dtSFM does not approximate a function: it parameterises the convergence equation that prescribes its architecture within the MRC framework^1,2^, and its size is set by the parameters of that equation rather than by the breadth of chemistry space it must cover. The model has approximately 42 million trainable parameters in total, trains end to end in approximately 15 hours on a single A100 GPU, and runs at inference on a single consumer-grade card; with drug and protein embeddings precomputed once and cached, each of the three applications completes in minutes-to-an-hour per run at cents-to-dollars per query. dtSFM was trained entirely on publicly available paired binding data, and the model weights, training pipeline, and embedding tools are released open-source. Substantial additional drug–target binding data, in both the continually growing public bioactivity databases and in private pharmaceutical collections, is not represented in the current corpus; any laboratory or company holding such data can augment the corpus and retrain or fine-tune the model on commodity hardware in days, then immediately deploy the updated model for their off-target, repurposing, or generative-design needs.

The results we report are entirely in silico. Every one of dtSFM’s three applications now requires wet-lab arbitration as the natural next step: chemoproteomic or kinase-panel confirmation of the predicted off-targets, biochemical or cellular validation of the repurposed and generated candidates against their intended targets, and orthogonal selectivity, ADMET, and pharmacological characterisation of the most promising candidates. AF3 verification is not a substitute for this experimental cycle. Several known limitations bound the present work: AF3 itself has documented blind spots (the hERG no-bind pattern we diagnose for the FLT3 series is one of these), the affinity head is preliminary, the decoder samples the chemistry distribution of its training data and does not by itself explore scaffold-novel space, and the published bioactivity corpus underrepresents many therapeutic target classes. dtSFM is a first SFM realisation at the encoder–decoder–design scale for one binding domain; its computational predictions are intended as a prioritised starting point for the experimental campaigns that will determine whether the predicted bindings are real.

## Methods

### Compute infrastructure

dtSFM encoder training, decoder training, encoder inference, and downstream evaluations were performed on Euler, ETH Zurich’s academic HPC cluster, with each job running on a single NVIDIA A100 40 GB GPU. AF3 cofolding at scale (more than 2,000 individual cofolds across the safety screening, repurposing, and generative-design protocols) was performed on the ALPS supercomputer at the Swiss National Supercomputing Centre (CSCS), whose compute nodes are built on NVIDIA Grace Hopper Superchip (GH200) modules pairing a Hopper-class H100 GPU with a Grace ARM CPU.

### Model architecture

#### Pre-trained encoders

The drug encoder is MoLFormer-XL^22^, a chemical language model pre-trained on approximately 1.1 × 10⁹ molecules, which produces a 768-dimensional global embedding from a canonical SMILES string. The protein encoder is ESM-2 650M ^24^, a protein language model, which produces a 1,280-dimensional per-residue representation from the target amino-acid sequence; protein sequences longer than 1,024 residues are truncated. Both encoders are loaded from their public Hugging Face checkpoints and held with weights frozen throughout dtSFM training.

#### Cross-attention encoder

The cross-attention encoder is a stack of two Transformer layers^19^ with eight attention heads per layer, model dimension d_model_ = 512, and feed-forward dimension d_ff_ = 2,048. The drug→protein and protein→drug cross-attention pathways are maintained as independent layer stacks to reflect the modality asymmetry between SMILES and protein sequence. Before cross-attention, the drug global embedding is projected from 768 to 512 dimensions by a single linear layer, per-atom drug features are constructed by an atom-level multilayer perceptron that takes element identity and Cartesian coordinates as input, and the protein per-residue features are projected from 1,280 to 512 dimensions and attention-pooled into a global protein representation. The cross-attention encoder contributes 14.4 million trainable parameters.

#### Output heads

Four heads read from the cross-attention encoder, each with a defined input, output, and loss function:

- **Global-retrieval head.** The retrieval head takes the drug and protein global features computed pre-cross-attention (each a 512-dimensional vector) and scores their binding compatibility as the cosine similarity of the two vectors in the shared 512-dimensional embedding space. Training uses the symmetric InfoNCE contrastive loss ^41^ with in-batch negatives and a learnable temperature τ.
- **Interface head.** The interface head takes the per-atom drug features computed post-cross-attention and outputs a per-atom logit for binding-interface membership. Training uses binary cross-entropy against per-atom interface labels derived from a 5 Å heavy-atom contact criterion.
- **Contact head.** The contact head takes the per-atom drug features and the per-residue protein features post-cross-attention, combines them bilinearly, and outputs a per-(atom, residue) contact logit. Training uses positively weighted binary cross-entropy to handle the high class imbalance, with a positive class rate of approximately 0.78%.
- **Affinity head.** The affinity head takes the pooled joint drug–target representation post-cross-attention and outputs a scalar regression on a unified *pK* scale that aggregates *pK_D_, pK_i_*, and *pIC_50_* values from the training corpus. Training uses mean squared error.

The four-head training loss is the sum of the individual losses weighted by α = (1, 1, 1, 0.5) for the retrieval, interface, contact, and affinity heads respectively, with per-head gradient magnitudes normalised by an exponential moving average so that contributions remain balanced across the course of training.

#### Decoder

The decoder is a cross-attentive autoregressive Transformer with model dimension d_dec_ = 512, structured as a decoder-only Transformer block with masked self-attention over the SMILES tokens generated so far and cross-attention over the projected protein features^19^. Tokenisation of SMILES uses the MoLFormer-XL tokenizer^22^ extended with an added end-of-sequence token, yielding a vocabulary of 2,363 tokens. The decoder is trained with teacher-forced cross-entropy on the ground-truth SMILES tokens of each (drug, protein) pair. The decoder follows the autoregressive language-model paradigm developed for biological-sequence generation^36,37^ and contributes approximately 27 million trainable parameters.

#### Total trainable parameters

The dtSFM encoder (cross-attention plus four output heads) and the decoder together contain approximately 42 million trainable parameters; the MoLFormer-XL and ESM-2 backbones are not updated.

### Training data and curation

#### PDBbind component

19,037 co-crystal protein–ligand complexes from PDBbind v2020 ^25^ with experimentally measured binding affinity (*K_D_, K_i_,* or *IC_50_* from the original deposition) were retained. Each entry contributes the protein amino-acid sequence, the canonical SMILES of the bound ligand, and a 5 Å heavy-atom contact map between the ligand atoms and protein residues derived from the deposited coordinates.

#### SAIR component

The bulk of the corpus is drawn from the Structurally Augmented IC50 Repository (SAIR) ^26^, which pairs bioactivities (*K_i_, IC_50_, K_D_*) curated from ChEMBL^27^ and BindingDB ^28^ with binding interfaces predicted by the Boltz-1x co-folding model ^11^. The SAIR release applies its own curation pipeline upstream: assays are restricted to direct binding measurements with standard relation’=’ and within a 1 pM to 100 μM dynamic range, ligands are filtered for PAINS and molecular weight ≤ 1,250 Da, proteins ≤ 2,000 residues are retained as monomeric assemblies, and any protein–ligand system with an experimentally solved structure already deposited in the PDB is excluded to prevent leakage against PDB-based training data. We applied an additional structural-confidence filter to the SAIR release, retaining only those entries with Boltz-1x iPTM ≥ 0.70 and overall confidence score ≥ 0.70. This filter produced 695,710 retained pairs, each supplying the same per-pair fields as PDBbind: protein sequence, canonical SMILES, pIC_50_, and a 5 Å heavy-atom contact map derived from the predicted complex.

#### Canonicalisation

All SMILES were canonicalised with RDKit^42^. Proteins were indexed by source-specific identifiers, *pdb:<PDB_ID>* for PDBbind and *uniprot:<UniProt_ID>* for SAIR, with sequences obtained from the deposited PDB or the corresponding UniProt entry. Because SAIR by construction excludes any protein–ligand system already present in the PDB, the two components contribute disjoint pairs to the combined corpus.

#### Combined corpus

The final curated corpus comprises 714,747 (drug, protein) pairs spanning 522,776 unique canonical SMILES and 22,964 unique protein sequences. Aggregated on the unified *pK* scale, affinities span *pK* = 0.40 to *pK* = 15.22 with a median of *pK* = 6.87.

#### Pre-computed embeddings

To avoid redundant encoder forward passes during training and inference, each unique drug was embedded once with MoLFormer-XL and each unique protein was embedded once with ESM-2 650M. The resulting drug embeddings (522,776 × 768 in half precision, ∼0.75 GB on disk) and protein embeddings (22,964 per-residue arrays of variable length × 1,280 in half precision, ∼25 GB on disk) were cached and read by the training loop rather than recomputed from sequence on each forward pass.

### Encoder training configuration

#### Optimisation

The cross-attention encoder and the four output heads were optimised end to end with AdamW ^43^ at learning rate 1 × 10⁻⁴ and weight decay 0.2, with a linear warmup over the first 5,000 training steps. Mixed-precision (bf16) training was used.

#### Batching and schedule

Batches of 128 (drug, protein) pairs were drawn at random from the 714,747-pair training partition. Training ran for 15 epochs at approximately 4,632 optimiser steps per epoch (∼70,000 total steps) on a single NVIDIA A100 40 GB GPU.

#### Multi-task loss balancing

The four output heads were summed with fixed weights α = (1, 1, 1, 0.5) for the retrieval, interface, contact, and affinity heads respectively, and each head’s gradient contribution was further normalised by an exponential moving average of its loss magnitude over the preceding training steps so that no single head dominated optimisation.

#### Validation and checkpointing

The held-out validation set described in the data splitting and leakage taxonomy section was evaluated at the end of every epoch, computing R@10 in both retrieval directions, interface and contact AUROC, and affinity Pearson r. A checkpoint was saved at each epoch (*epoch_001.pt* through *epoch_015.pt*). The locked checkpoint used for every downstream result in this paper is *epoch_010.pt* of training run *b3_20260506_191149*, the earliest checkpoint at which both retrieval directions had reached their validation plateau (**Fig. 2c**).

### Decoder training configuration

#### Tokenisation

The decoder uses the MoLFormer-XL tokenizer^22^ extended with an added end-of-sequence token, yielding a vocabulary of 2,363 tokens including the standard *<BOS>*, *<EOS>*, and *<PAD>* symbols. SMILES are encoded with this tokenizer for both teacher-forced training and sampling.

#### Optimisation

The decoder was optimised with AdamW ^43^ at peak learning rate 2 × 10⁻⁴ with cosine decay to a minimum of 1 × 10⁻⁶ over the training schedule, with a linear warmup over the first 2,000 training steps. Mixed-precision (bf16) training was used.

#### Batching and schedule

Batches of 64 (target_protein_embedding, ground_truth_SMILES) pairs were drawn from the same 714,747-pair training partition as the encoder. SMILES were truncated to 256 tokens and protein sequences to 1,024 residues. Training ran for 100,000 optimiser steps on a single NVIDIA A100 40 GB GPU. The locked checkpoint used in this paper is the model state at training step 50,000 (*decoder_v0.2_step50K*).

#### Loss

The decoder was trained with teacher-forced cross-entropy on the ground-truth SMILES tokens of each (drug, protein) pair.

### AlphaFold-3 cofolding protocol

#### Deployment

AF3 was deployed on ALPS via the official AF3 container packaged as a Container Engine for Application Containers (EDF) image, and was run with SLURM job arrays of up to several hundred cofolds in parallel.

#### Input construction

Each cofold input is a JSON specification containing the target protein sequence with its precomputed MMseqs2-derived multiple sequence alignment injected verbatim as the *unpairedMsa* field, the candidate compound SMILES as the corresponding ligand entity, an empty *pairedMsa* field, and an empty *templates* list. The multiple sequence alignment is generated once per unique target sequence with the MMseqs2 ^29^ pipeline of ColabFold^44^ against the UniRef30 sequence database and the ColabFoldDB metagenomic database, then converted to A3M format and cleaned of characters AF3 rejects before injection. The pre-generated MSAs are produced on Euler and copied to ALPS before each cofolding job, eliminating online sequence-database searches at inference time.

#### Sampling

AF3 was run with *modelSeeds = [42]* and the default of five diffusion samples per seed, yielding five candidate structures per cofold input.

#### Output processing

For each cofold, the resulting model structure (CIF format) and the corresponding summary-confidence and per-residue-confidence JSON files were retained. The interface predicted TM-score (iPTM), the interface predicted aligned error (PAE, taken as the minimum interface-PAE value across heavy-atom interface contacts), and the per-chain-pair interface PTM were extracted from the summary-confidence JSON for downstream analyses. Each cofold was scored against the binder gate (iPTM ≥ 0.7 and interface PAE ≤ 5 Å) and the anchor-grade gate (iPTM ≥ 0.9 and interface PAE ≤ 1.67 Å) defined in **Fig. 6c**.

### Data splitting and leakage taxonomy

#### Protein-cluster split

The 22,964 unique training proteins were clustered by amino-acid sequence identity with MMseqs2 ^29^ at the 80% identity threshold (*--min-seq-id 0.8 --cov-mode 0-c 0.8*). Entire clusters were assigned to a single partition so that no validation or test protein shares more than 80% sequence identity with any training protein. The resulting partition holds 592,888 (drug, protein) pairs in the training set, 65,951 pairs in the validation set, and 55,908 pairs in the test set, summing to the full 714,747-pair corpus. The validation and test sets each cover 2,296 unique proteins in 2,296 unique clusters.

#### Drug-novelty (A/B/C) taxonomy

Independently of the protein-cluster split, each evaluated (drug, protein) pair is graded along a drug-novelty axis. The pair is Class A if the identical (drug, protein) pair was seen in training (memorisation); Class B if the drug was seen in training but never paired with this protein (drug-level generalisation against a held-out protein); and Class C if the drug is novel to training (chemistry novelty). The protein-cluster holdout removes any Class A pair from the validation and test sets by construction, so the validation split partitions into 24,753 Class B pairs (38%) and 41,198 Class C pairs (62%).

#### Direct leakage verification

The split was verified by direct pairwise overlap measurement between training and each held-out set across four dimensions: exact (drug, protein) pair identity, drug identity by canonical SMILES, protein identity, and MMseqs2 cluster identity. The full verification is reported in the Leakage verification section.

#### Axis usage throughout the paper

“In-distribution” (ID) and “out-of-distribution” (OOD) refer to the protein-cluster axis throughout the paper, distinguishing pairs whose protein belongs to a training cluster from pairs whose protein belongs to a held-out cluster. “Class A/B/C” refers to the drug-novelty axis. The encoder evaluation in **Fig. 3** uses the protein axis; the application protocols in **Fig. 4** (off-target safety screening) and **Fig. 5** (library repurposing) use the drug axis.

### Encoder evaluation

#### Sampling

For each of three evaluation regimes (in-distribution, where the protein cluster appears in the training set; out-of-distribution validation, where the protein cluster is in the validation partition; and out-of-distribution test, where the protein cluster is in the test partition), 4,000 (drug, protein) pairs were sampled uniformly from the corresponding partition with a fixed random seed.

#### Retrieval

Retrieval was evaluated full-pool and bidirectionally. In the drug→target direction, the true protein was ranked against the complete set of unique proteins represented in the sample, with a pool of 928 proteins for in-distribution, 311 for out-of-distribution validation, and 349 for out-of-distribution test. In the target→drug direction, the true drug was ranked against the complete set of unique drugs in the sample, with a pool of approximately 3,950 drugs in every regime. Recall at rank K (R@K) was computed for K ∈ {1, 5, 10, 50, 100} in both directions, and reported with 95% confidence intervals from 1,000 bootstrap resamples of the query set.

#### Structural heads

The interface head and the contact head were evaluated as binary classifiers by the area under the receiver-operating curve (AUROC), computed over all per-atom calls for the interface head and over all per-(atom, residue) cells for the contact head; chance corresponds to AUROC 0.5.

#### Affinity head

The affinity head was evaluated by Pearson correlation r between its predicted pK and the training-target pK on each evaluated pair, reported per regime.

### Leakage verification

#### Direct pairwise overlap

For both the validation and test sets, overlap with the training set was measured directly across four dimensions: (i) exact (drug, protein) pair identity; (ii) drug identity by canonical SMILES; (iii) protein identity by exact sequence match; and (iv) MMseqs2 cluster identity. The validation and test sets contained zero exact pair overlaps with training (0 of 65,951 and 0 of 55,908 pairs respectively), zero protein overlaps (0 of 2,296 unique proteins each), and zero MMseqs2-cluster overlaps (0 of 2,296 unique clusters each). Drug overlap by canonical SMILES was 36% for validation and 37% for test, reflecting the prevalence of compounds tested against multiple protein targets in the public corpus; this overlap defines the Class B versus Class C split along the drug-novelty axis described in the Data splitting and leakage taxonomy section.

#### Chemistry-novelty characterisation

The chemical novelty of each held-out drug relative to the training corpus was characterised by the maximum ECFP4 Tanimoto similarity (radius 2, 1,024 bits) ^45^ of the held-out drug to any drug in the training set, computed on a 5,000-drug uniform random subsample of each held-out split. Across both splits, the median maximum Tanimoto was approximately 0.69. Approximately 38% of held-out drugs were near-duplicates of a training drug (Tanimoto 0.9–1.0), 12% were close analogues (0.7–0.9), 21% were moderately similar (0.5–0.7), 28% fell in the novel-scaffold range (0.3–0.5), and 1–2% had no close training analogue (Tanimoto < 0.3).

### Off-target safety screening protocol

#### Gene-rank computation

For each query drug, the dtSFM encoder computed the retrieval cosine between the drug and each of the 22,964 training proteins using the locked *epoch_010* checkpoint. Multiple training-protein entries belonging to the same gene (across PDB structures, paralogues, or species variants) were collapsed to a single gene-level score by taking the maximum cosine across that gene’s protein entries, yielding a ranked list of 4,910 unique training genes per query drug.

#### Reference dataset

The kinome-wide off-target landscape used for validation was the Klaeger et al. chemoproteomic kinobeads dataset^5^. Nine clinical kinase inhibitors were analysed: the in-training inhibitors dasatinib, erlotinib, gefitinib, imatinib, sorafenib, and sunitinib, and the drug-out-of-distribution inhibitors crizotinib, ibrutinib, and ponatinib (each with its canonical SMILES absent from the training corpus). The kinases common to all nine inhibitors in the Klaeger data formed a 276-kinase comparison set, and binders were defined by the Klaeger-reported Kd ≤ 30 µM threshold.

#### Class B no-leakage recovery

For the leakage-controlled results, the documented off-target pairs were restricted to Class B, namely pairs in which both the drug and the gene appeared in the training corpus but never as this specific pair. This restriction yielded 27 Class B off-target pairs across five inhibitors (dasatinib, erlotinib, gefitinib, imatinib, sunitinib). For each Class B pair, the gene rank of the off-target in the inhibitor’s dtSFM screen was recorded, and cumulative recovery at top-10, top-50, top-100, and top-500 of the 4,910-gene screen was reported.

#### Per-drug separation AUROC

For each of the nine inhibitors, the AUROC was computed for the dtSFM gene-rank-based separation of the inhibitor’s Klaeger-measured binders from its measured non-binders, restricted to the 276-kinase comparison set.

### Library repurposing screening protocol

#### Library screen

For each target, the dtSFM encoder computed the retrieval cosine between the target sequence and each of the 522,776 unique compounds in the training library using the locked *epoch_010* checkpoint, producing a complete ranked list of the library in a single pass per target. Drugs and proteins were embedded once and cached (see Training data and curation), so each per-target screen amounted to a single ESM-2 forward pass for the target plus 522,776 cross-attention forward passes for the (target, drug) pairs.

#### Target selection

Three clinically relevant immunology proteins were screened: NLRP3, CD73, and STING1. These targets were chosen to span the leakage-class spectrum of training-set coverage: CD73 is Class A–dominant in its top hits (heavily represented in training); STING1 is Class B–dominant (its canonical clinical agonists are absent from the library and therefore all Class C); and NLRP3 sits between them.

#### Anchor and negative-control compounds

A set of published anchor compounds was assembled for each target from the clinical and chemical-biology literature, comprising the canonical inhibitor or agonist, additional clinical-stage compounds, and substrate or natural-ligand analogues where applicable; each anchor was labelled by its leakage class (Class A, B, or C). Five negative-control compounds per target (approved drugs not known to bind the target: Loratadine, Amoxicillin, Levothyroxine, Warfarin, and one further per-target control) were included as decoy entries.

#### Verification cohort sampling

From each target’s ranked screen, a verification cohort was assembled by combining the top-50 cosine hits, a scaffold-diverse sample of 35–50 next-tier hits (selected to maximise ECFP4 Tanimoto-diverse coverage of the high-cosine region), the published anchor compounds for the target, and the five matched negative controls. The per-target totals were 95 pairs for CD73, 115 for NLRP3, and 113 for STING1, summing to 323 pairs in total. Each cohort pair was cofolded with AF3 as described above.

#### Off-library anchor synthetic ranking

Published anchor compounds that were not present in the v3 training library (notably all canonical STING1 agonists, which are Class C) were scored by encoding the anchor’s SMILES with MoLFormer-XL and computing the cross-attention cosine against the target embedding; the resulting cosine was inserted into the library cosine distribution to produce a synthetic rank within the 522,776-compound library. The same procedure was applied to the three Class C complementarity-illustration compounds of **Fig. 5a** (Dapansutrile for NLRP3; MSA-2 and diABZI compound 3 for STING1).

#### Novel-candidate definition

A novel candidate was defined as a (compound, target) pair drawn from the dtSFM-prioritised strata (top-50 cosine or scaffold-diverse) that is Class B or Class C and that satisfies the AF3 binder gate (iPTM ≥ 0.7 AND interface PAE ≤ 5 Å, as defined in the AlphaFold-3 cofold protocol and **Fig. 6c**).

### Generative design protocol

#### Target selection

Sixteen clinically relevant immunology and oncology proteins were selected for generative design: ALK, BTK, CDK4, CDK6, EGFR, F11, FLT3, JAK1, JAK2, JAK3, MAP2K1, MAP2K2, PARP1, PARP2, PIK3CA, and TYK2.

#### Decoder sampling

For each target, the cross-attentive decoder (checkpoint *decoder_v0.2_step50K*) generated approximately 75 candidate SMILES strings by temperature sampling (temperature 0.8, maximum 100 new tokens per sample), conditioned on the per-residue protein features of the target produced by ESM-2. Each generated token sequence was parsed with RDKit, and samples that failed to parse to a valid molecule were discarded. The combined generation across the 16 targets produced 1,200 valid candidate molecules.

#### Encoder reranking and drug-likeness filter

The 1,200 valid candidates were reranked per target by the dtSFM encoder cosine between the candidate’s encoded SMILES and the target’s encoded sequence, and filtered by basic drug-likeness criteria (QED and Lipinski Ro5), yielding 849 reranked candidates retained for chemistry-novelty characterisation.

#### AlphaFold 3 cofolding

All 1,200 generated candidates (the full generation cohort, not the reranked subset) were cofolded with AF3 against their intended target following the AF3 cofold protocol above. In addition, an approved-drug anchor for each target was cofolded against the same target as a positive reference compound, yielding 15 approved-drug anchor cofolds across the 16 targets. Eighty negative-control decoy drugs (five per target, comprising approved drugs not known to bind the respective target, such as Loratadine, Amoxicillin, Levothyroxine, Warfarin, and one further per-target control) were also cofolded as a non-binder baseline.

#### Confidence gates

Two AF3 confidence gates were used to score the cofolds. The broad binder gate (interface predicted TM-score iPTM ≥ 0.7 AND interface predicted aligned error PAE ≤ 5 Å) is the same gate used in the library repurposing protocol. The stricter anchor-grade gate (iPTM ≥ 0.9 AND interface PAE ≤ 1.67 Å) has a PAE bound equal to the worst PAE observed among the 15 approved-drug anchor cofolds, so that all 15 anchors satisfy it by construction; a generated candidate that clears the same gate is therefore as confident a binder by AF3 as the approved drug for that target.

#### Chemistry novelty characterisation

The chemistry novelty of the reranked cohort was characterised by two ECFP4 (radius 2, 1,024-bit) ^45^ Tanimoto similarity measures: the maximum Tanimoto of each candidate to its nearest compound in the 522,776-compound training library (median 0.73, characterising novelty relative to the training distribution), and the maximum Tanimoto of each candidate to its target’s approved drug (median 0.22, characterising novelty relative to the approved chemistry for the target).

### Orthogonal AI verification

#### Structural verification

Every structural prediction in this work was checked by AF3 as an orthogonal verifier. AF3 shares no architecture, training data, or learned representation with dtSFM, and on the candidate sets the dtSFM encoder cosine is essentially uncorrelated with AlphaFold-3 confidence (Pearson r ≈ 0); structural agreement therefore constitutes independent corroboration rather than circular confirmation.

#### Coding Audit

Every quantitative claim in this paper was independently re-derived by a second AI coding assistant (Codex, OpenAI) acting as an orthogonal auditor. Operating in a clean session with access only to the artifacts committed to the public repository — and no access to the development sessions that produced them — the auditor recomputed each claim from its source data and graded it pass, partial, or fail. The audit closed at 48 pass / 3 partial / 0 fail, with no numerical claim found to be in error (the three partials are documented bookkeeping items); the four direct leakage measurements (exact drug–protein pair, drug identity, protein cluster, and Tanimoto bucket) reproduced the reported values. The complete audit record, the claim-to-file map, and all verification scripts are provided in the code repository (github.com/Reddy-BIIE-ETHZ/dtSFM, audit/) and the archived data deposit (Zenodo, DOI 10.5281/zenodo.20581780).

### Vibe Coding Starter Prompt

#### Applying dtSFM to your target or library

dtSFM is designed for direct use by experimental biologists with no coding background. The block below is a single prompt: paste it into an AI coding agent, such as Claude Code (Anthropic) along with the published dtSFM PDF. Claude Code will pull the repository and model weights, install the environment, and walk you through your specific use case in plain language. The only fields you must fill in are your name and the use-case checkbox; every other field is optional — Claude Code can fetch the target sequence from UniProt, choose default hyperparameters, and identify the closest approved binder by itself when those fields are left blank.

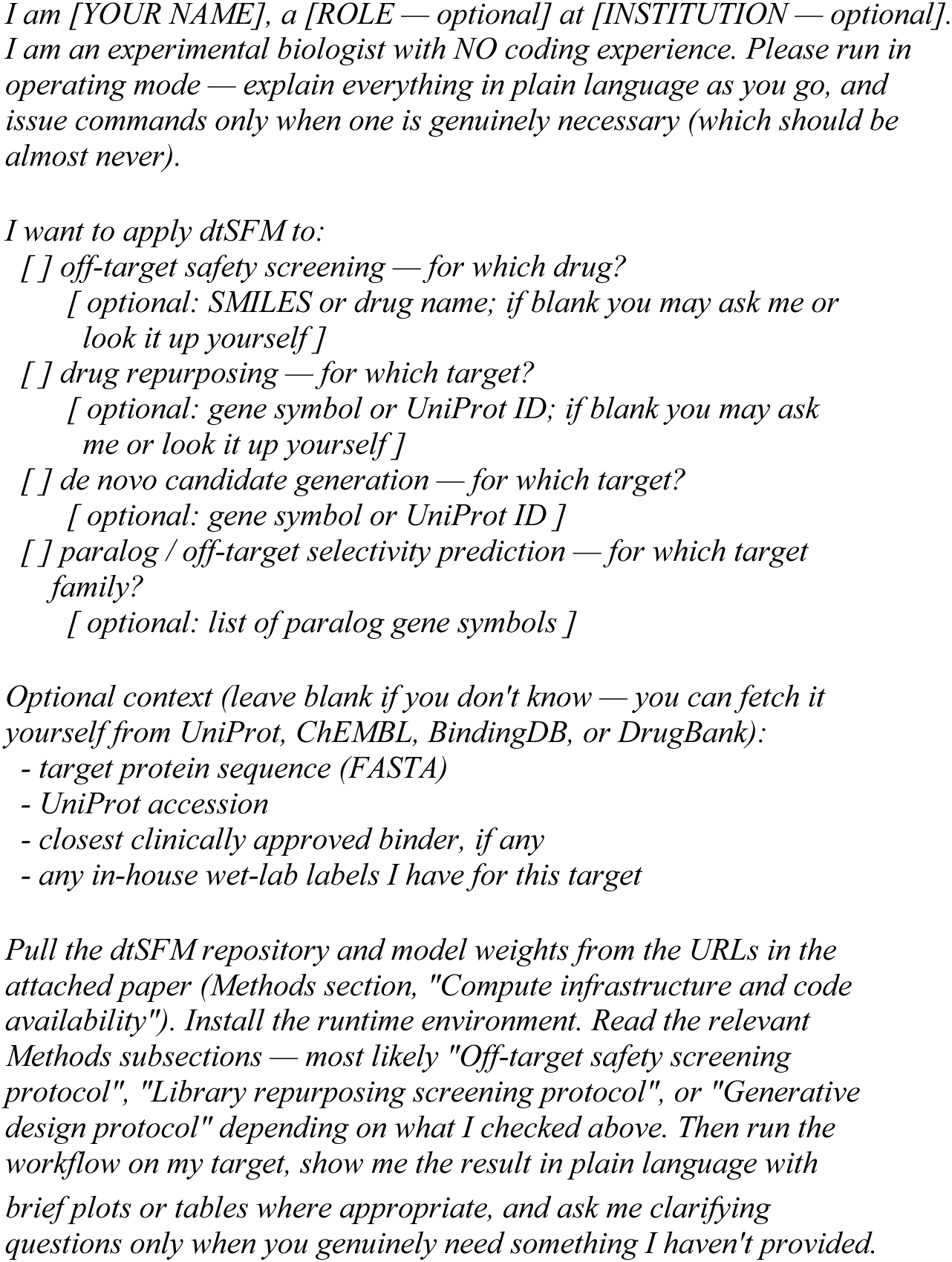

## Acknowledgments

ETH Zurich Scientific IT Services for High Performance Computing and Swiss National Supercomputing Centre (CSCS) provided computing resources and excellent support. Thank you to Riyaz Khan and Roy Ehling for testing the “Vibe Coding Drug Discovery” protocol and additional feedback.

## Data and code availability

All training data were derived from publicly available databases (see **Table 1**). dtSFM has approximately 42 million trainable parameters — a cross-attention encoder paired with a generative decoder — and can be trained in approximately 15 hours on a single GPU. Pre-trained encoder and decoder weights are available on Hugging Face: https://huggingface.co/SFM-BIIE-ETHZ/dtSFM-v3. Code, training and evaluation pipelines, the three application pipelines, and orthogonal audit records are available on GitHub: https://github.com/Reddy-BIIE-ETHZ/dtSFM. Generated candidate pools and AlphaFold-3 cofold structures are deposited on Zenodo: https://doi.org/10.5281/zenodo.20581780.

## Supplementary Material

### Contents

**Figure S1.** Direct leakage audit: pair / drug / protein / MMseqs2-cluster overlap between training and each held-out split.

**Figure S2.** Chemistry-novelty distribution: ECFP4 Tanimoto of held-out drugs to nearest training drug.

**Figure S3.** Affinity head — preliminary: predicted vs measured pK regression, ID vs OOD.

**Figure S4.** Decoder base sampling: QED, Lipinski-Ro5 per target, encoder cosine, chemistry novelty.

**Figure S5.** Decoy vs design AlphaFold-3 gate-pass rates.

**Figure S6.** Structural wall: 149 individual design–drug cofolds (top-10 per target). Due to file size, the complete 17-page gallery (all 16 targets × up to 10 designs, anchor-overlaid, with per-design AlphaFold-3 iPTM/PAE) is hosted at the project repository: https://github.com/Reddy-BIIE-ETHZ/dtSFM/blob/main/supplementary/figS6_structural_gallery.pdf

**Table S1.** SMILES and computed properties of all generated designs shown in the structural-validation gallery (Fig. 7), with the approved-drug anchor for each of the 16 targets.

---

**Figure S1.**
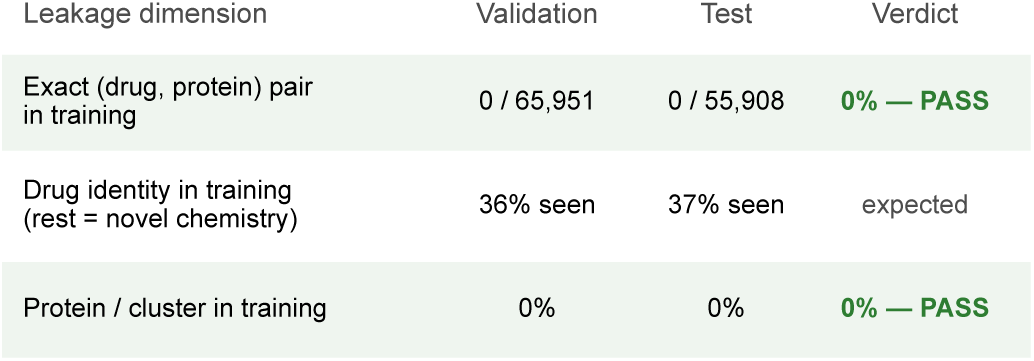
Direct leakage audit across the train/validation/test split. Pairwise overlap between the training set and each held-out split along three dimensions. Exact (drug, protein) pair overlap was zero in both validation (0 / 65,951) and test (0 / 55,908); protein-identity and MMseqs2-cluster overlap were zero (the split holds out entire sequence clusters); drug-identity overlap by canonical SMILES was 36% (validation) and 37% (test), which is expected and defines the Class B / Class C split rather than pair leakage.

**Figure S2.**
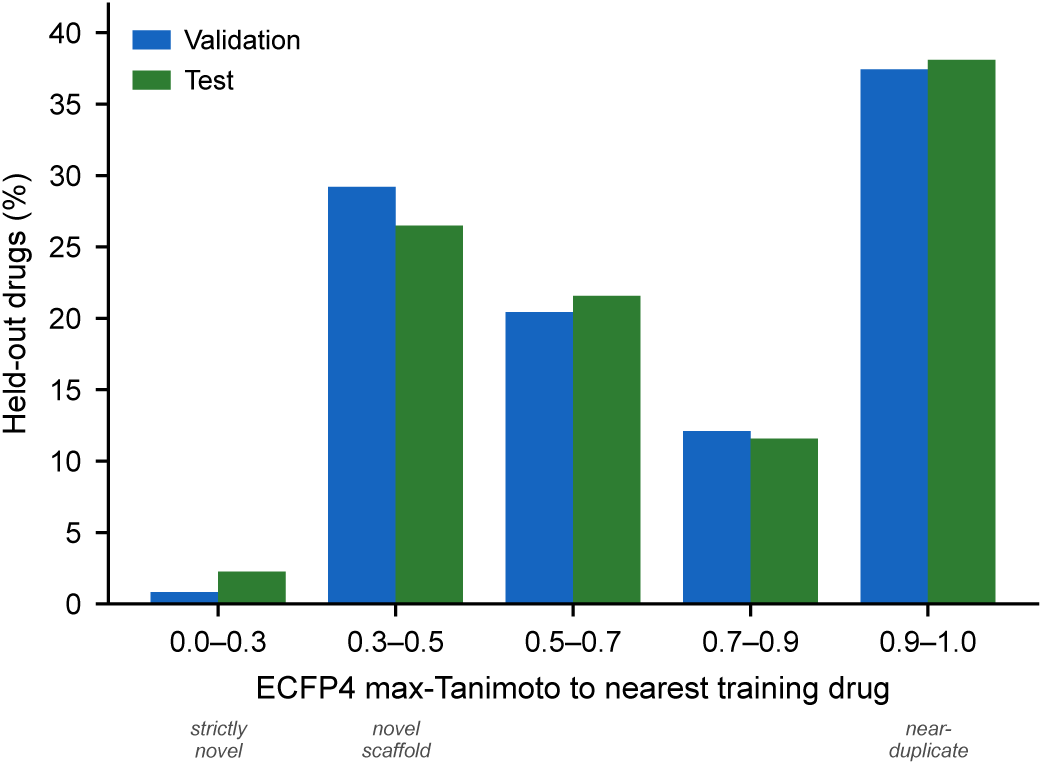
Chemistry-novelty distribution of held-out drugs relative to training. Maximum ECFP4 Tanimoto similarity (radius 2, 1,024-bit) of each held-out drug to its nearest training-set drug, on a 5,000-drug random subsample of each split (validation, test; seed 42), binned into five ranges. About 37% of held-out drugs were near-duplicates of a training compound (Tanimoto 0.9–1.0), ∼29% (validation) / ∼26% (test) fell in the 0.3–0.5 novel-scaffold range, and only ∼0.8% / ∼2.3% were strictly novel (< 0.3); median maximum Tanimoto ≈ 0.69. The Class C cohort is thus novel by exact-SMILES identity but consists largely of scaffold variants of training chemistry rather than wholly unprecedented structures.

**Figure S3.**
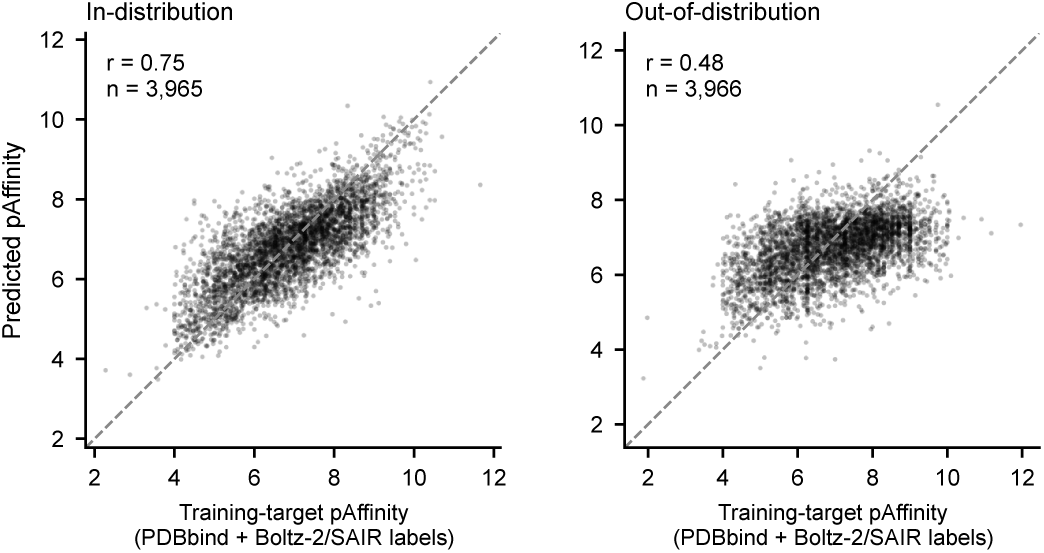
Affinity head — preliminary measured-affinity performance. Predicted versus measured pK for the affinity head, in-distribution and out-of-distribution. The head was trained on PDBbind and SAIR labels (real measured *Kd / IC50 / EC50* from ChEMBL and BindingDB) and reproduced the held-out measured label at Pearson r = 0.75 (ID) / 0.48 (OOD validation) / 0.37 (OOD test).

**Figure S4.**
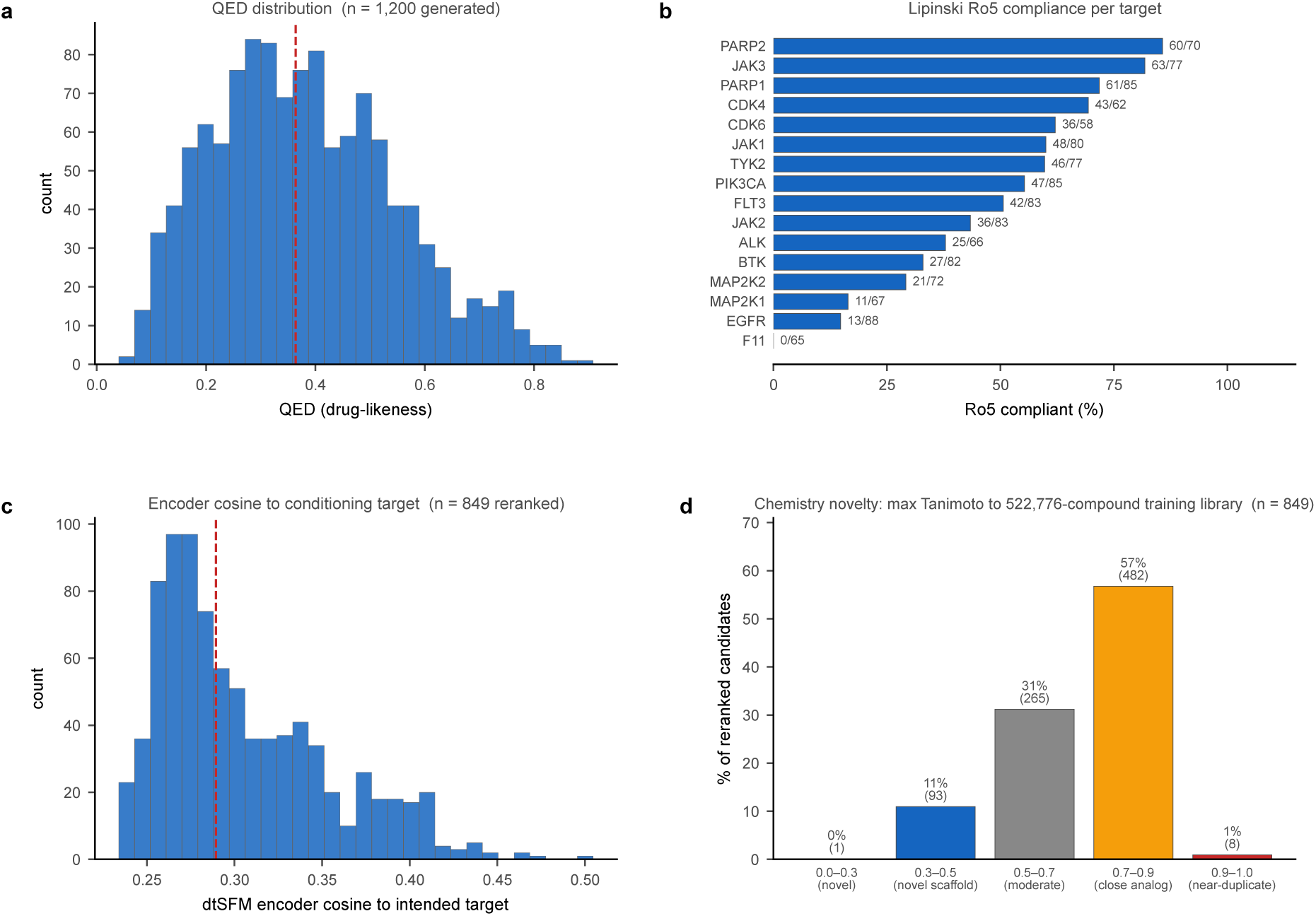
Base sampling performance of the cross-attentive decoder. Characterisation of decoder output across the 16 targets. (**a**) QED distribution of the 1,200-candidate cohort (median 0.36). (**b**) Lipinski Ro5 compliance per target. (**c**) dtSFM-encoder cosine of each reranked generated molecule to its intended target (n = 849), confirming the decoder samples molecules the encoder recognises as target-compatible. (**d**) Chemistry novelty: maximum ECFP4 Tanimoto of each reranked candidate to its nearest compound in the 522,776-compound training library (median 0.73; none a duplicate).

**Figure S5.**
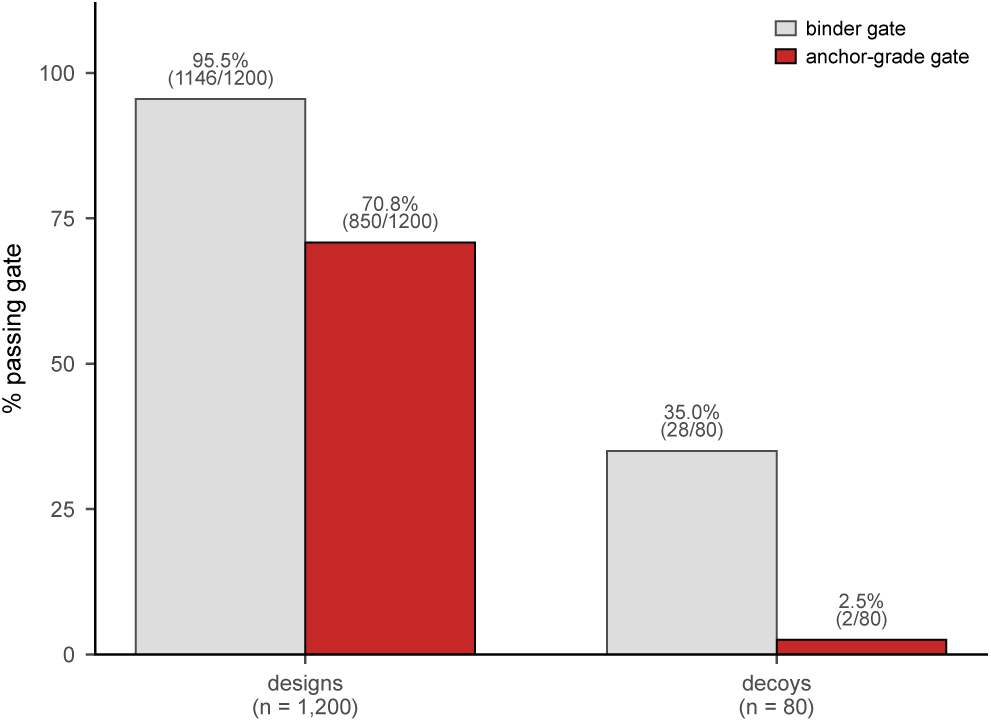
Negative-control decoys are excluded by the AF3 anchor-grade gate. Fraction of generated designs versus negative-control decoy drugs clearing each AF3 gate. Of the 1,200 designs, 1,146 (95.5%) cleared the binder gate (iPTM ≥ 0.7 AND interface PAE ≤ 5 Å) and 850 (70.8%) the anchor-grade gate (iPTM ≥ 0.9 AND interface PAE ≤ 1.67 Å); of 80 decoys, 28 (35%) cleared the binder gate but only 2 (2.5%) the anchor-grade gate. The binder gate sits at the AF3 drug-like floor, whereas the anchor-grade gate effectively separates designs from decoys.

**Figure S6.**
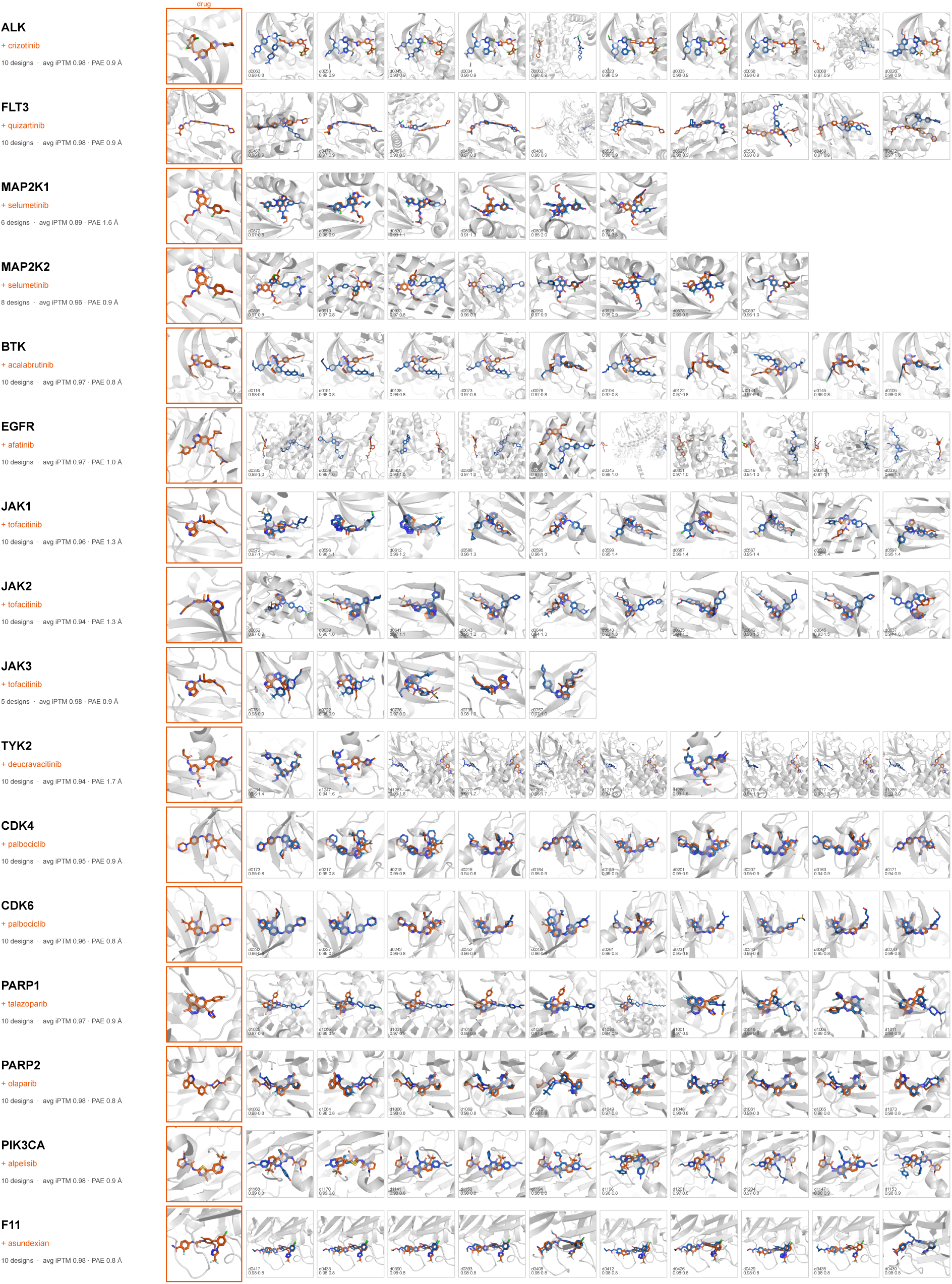
Structural-validation wall: every top design overlaid on the approved drug. The exhaustive companion to Fig. 7. For each of the 16 targets, every top-10 AlphaFold-3-confident generated design (blue) is individually overlaid on the approved drug (orange) in the target pocket (149 single design–drug cofolds in total), each tile annotated with the design identifier and its iPTM / interface PAE; row labels give the target, approved drug, and ensemble-mean confidence.. Full structural gallery file available here: https://github.com/Reddy-BIIE-ETHZ/dtSFM/blob/main/supplementary/figS6_structural_gallery.pdf

**Table S1.**
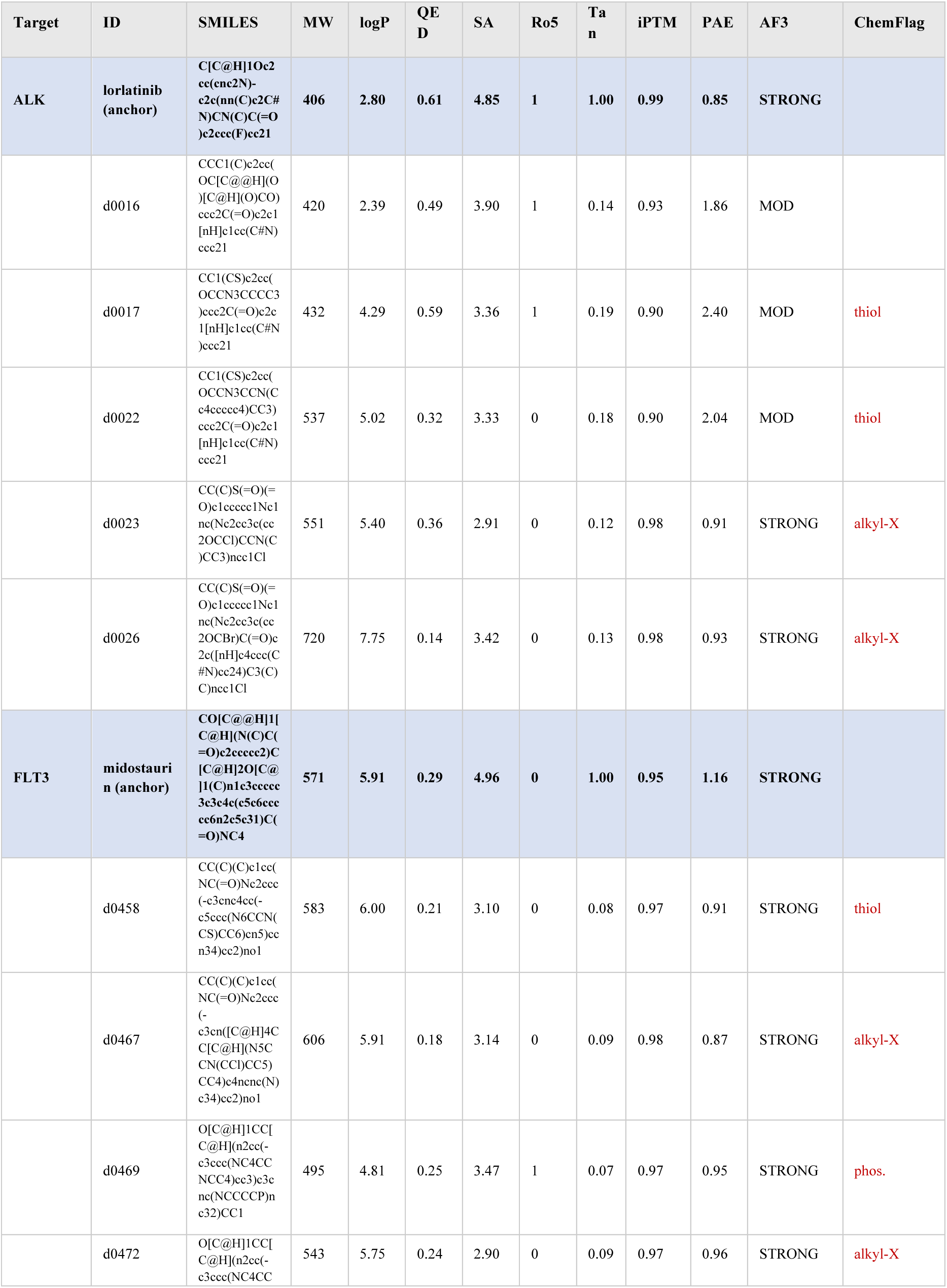
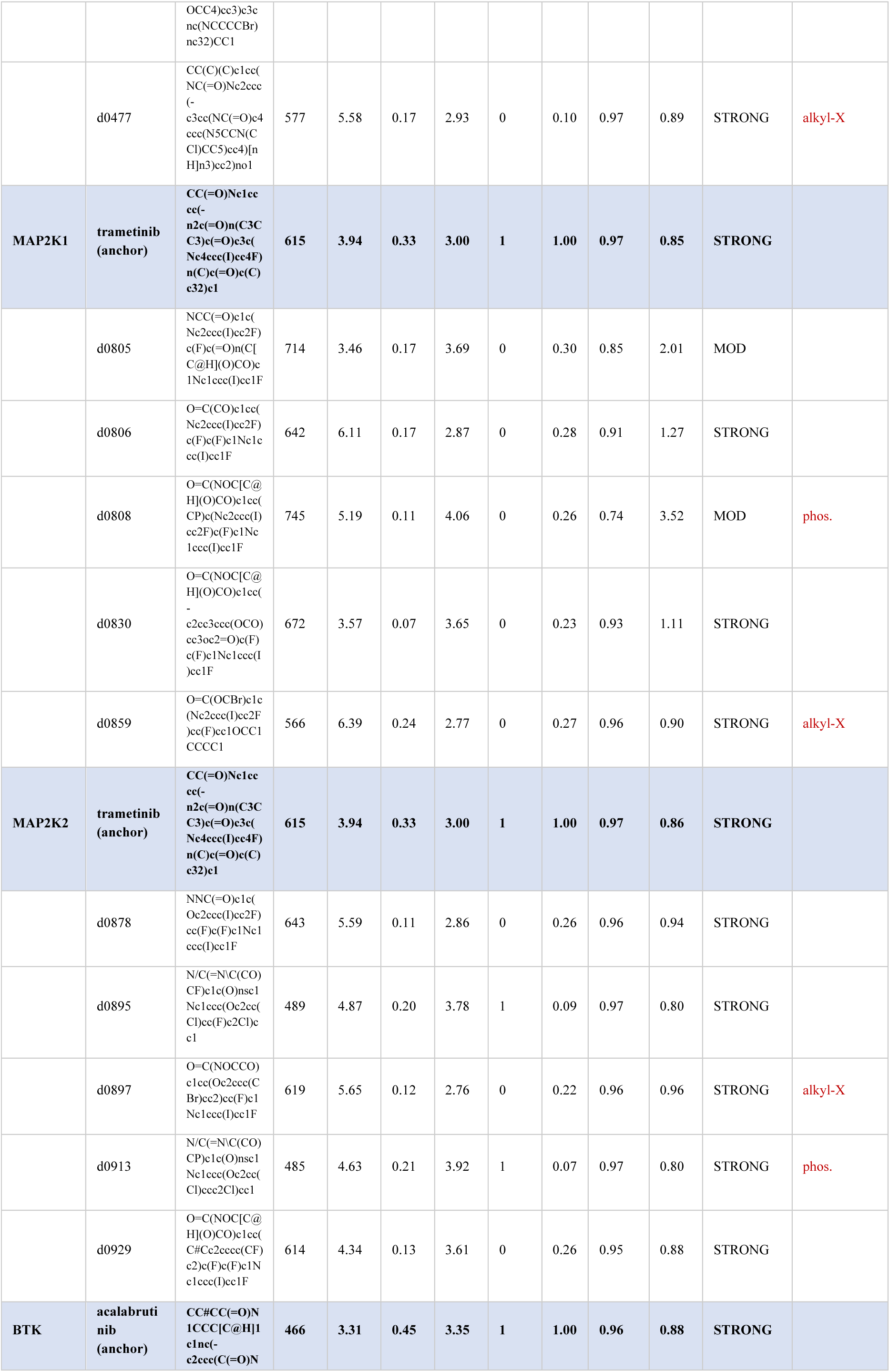
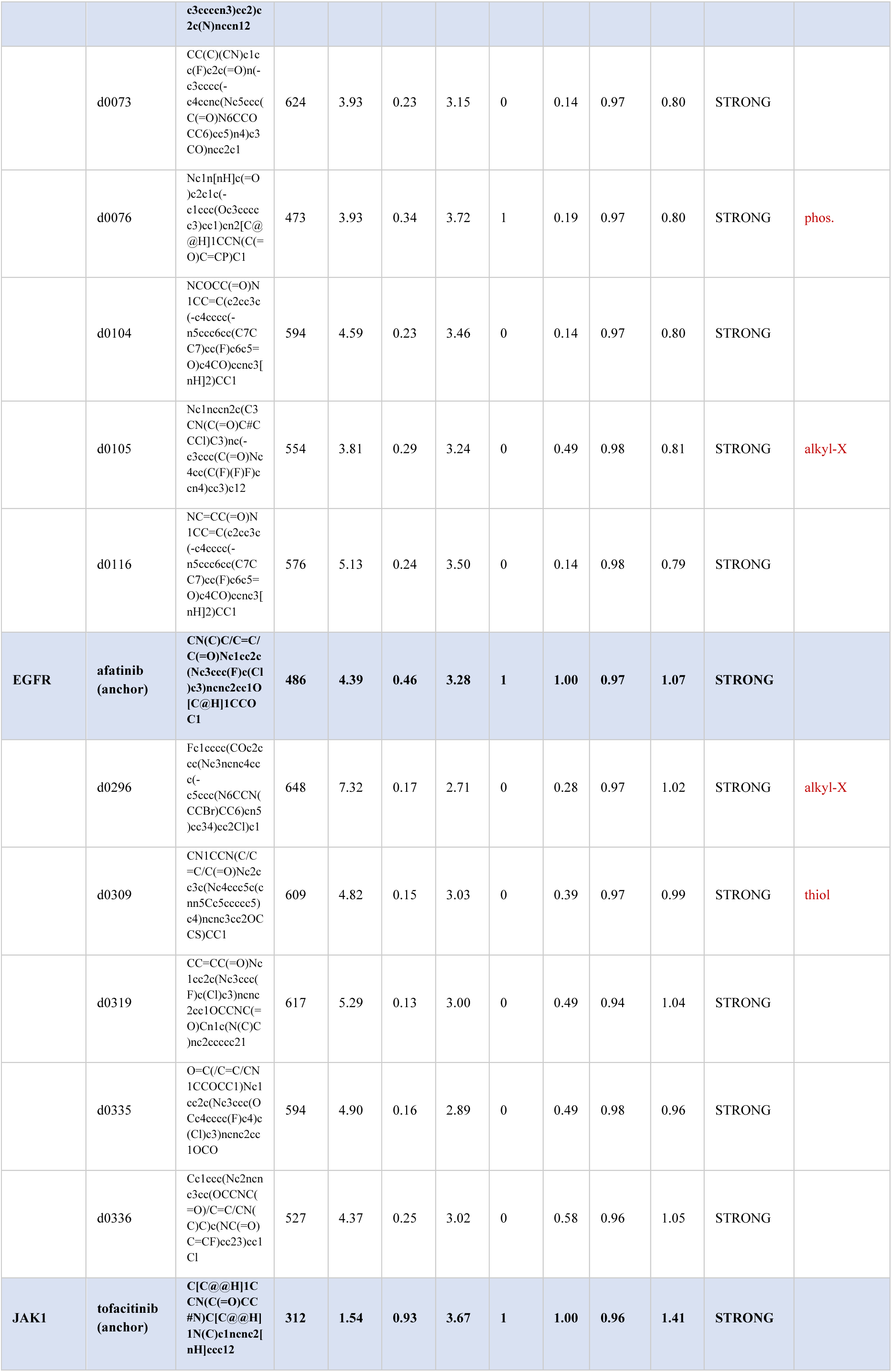
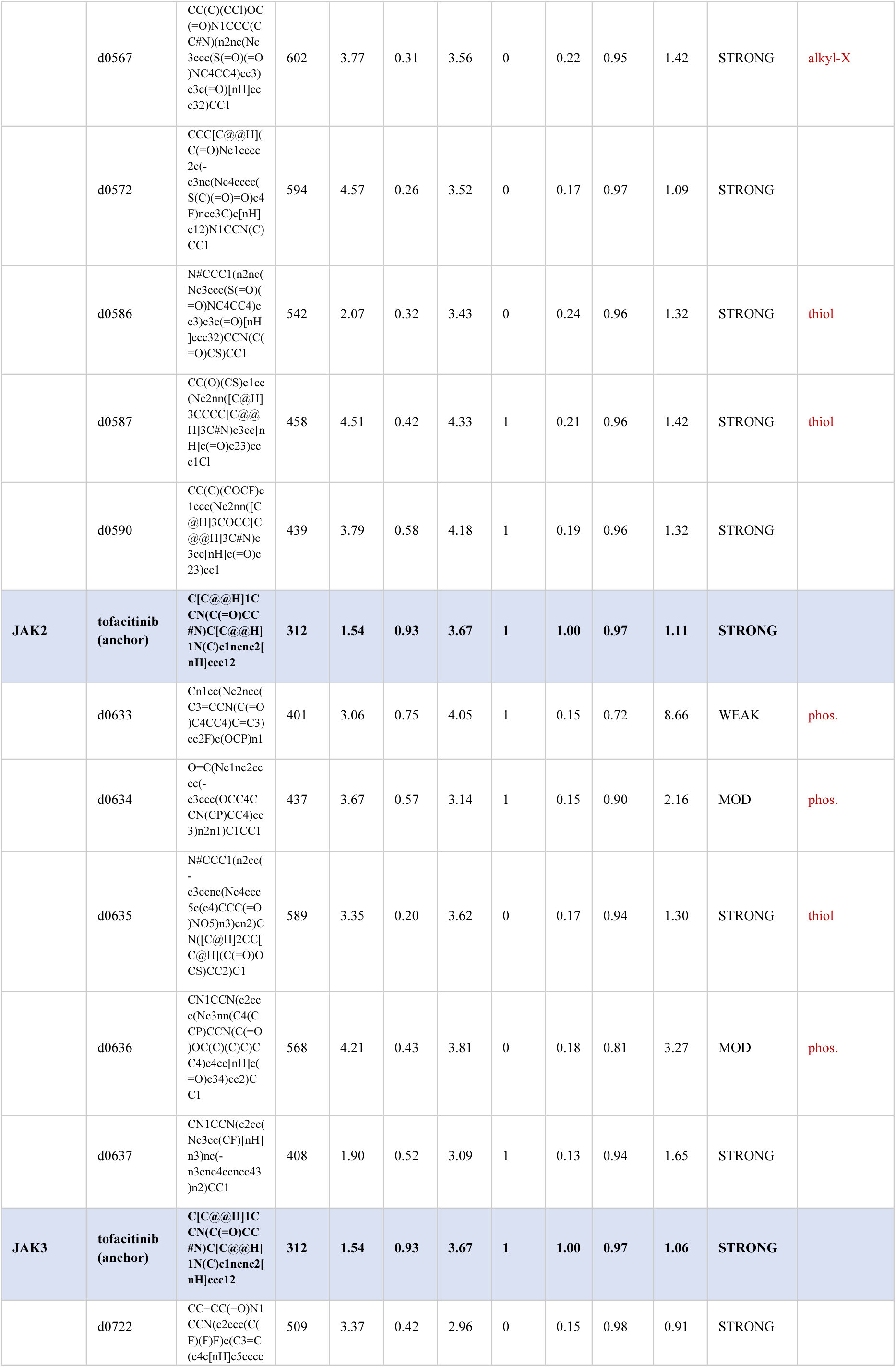
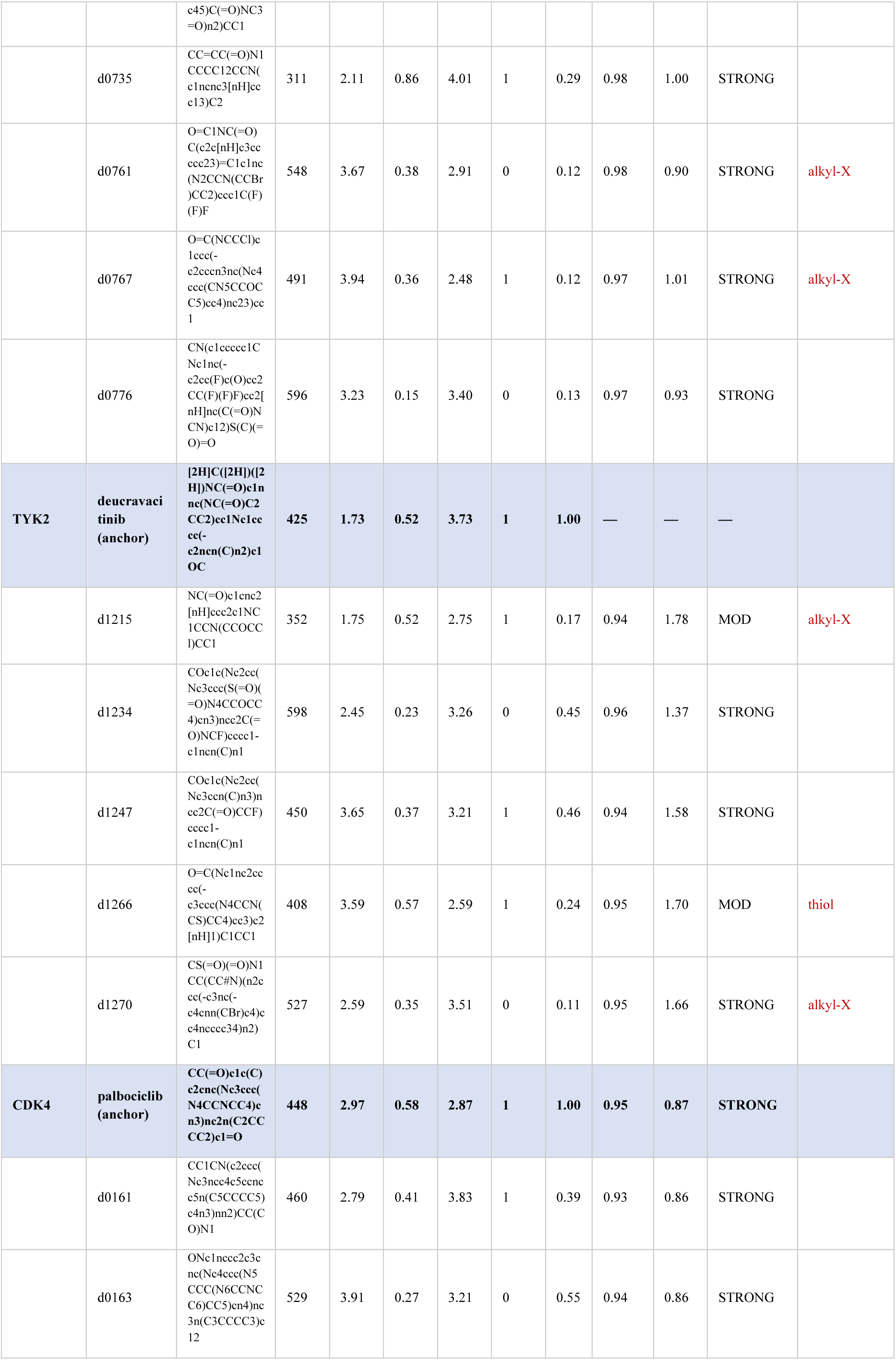
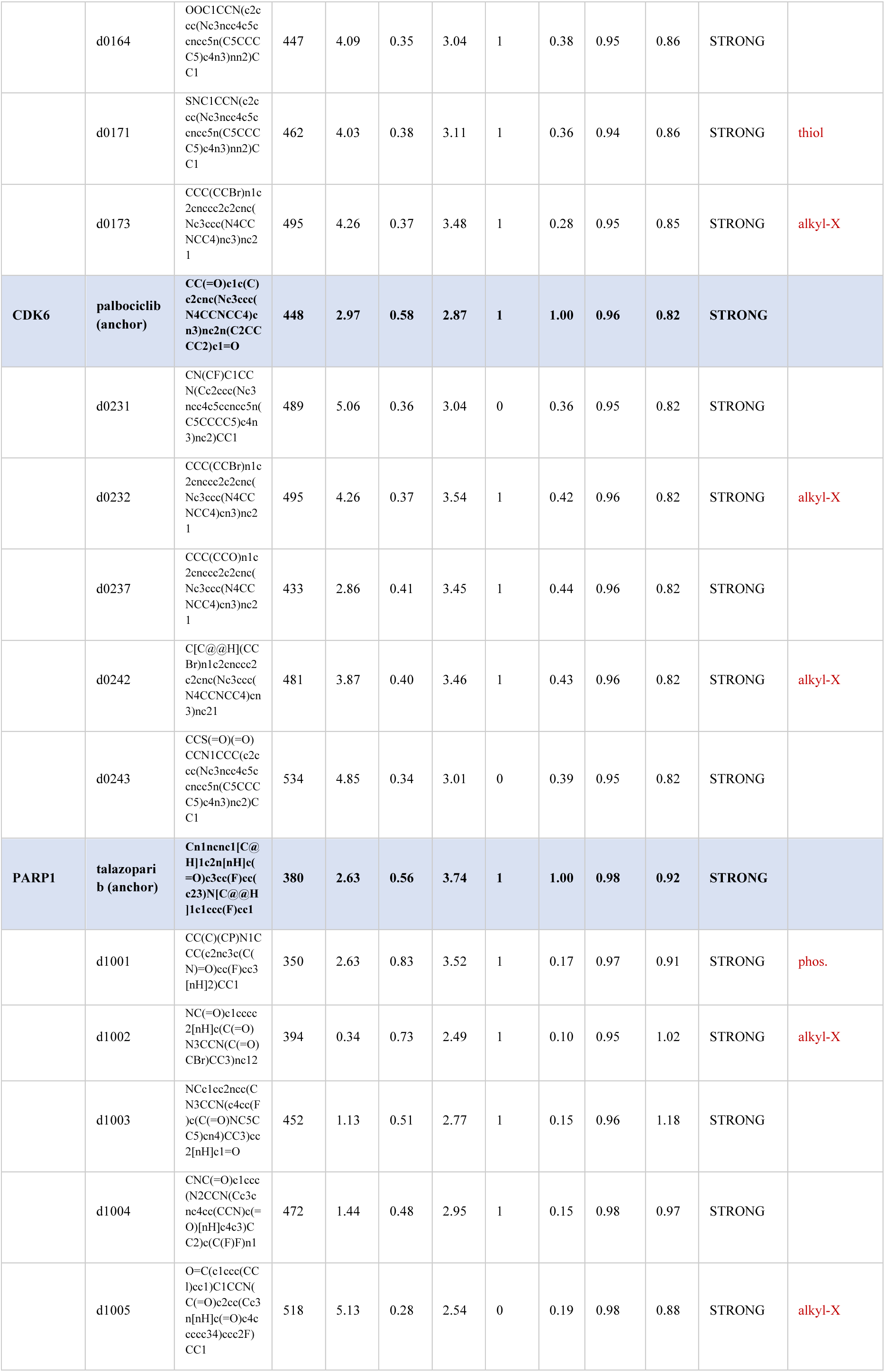
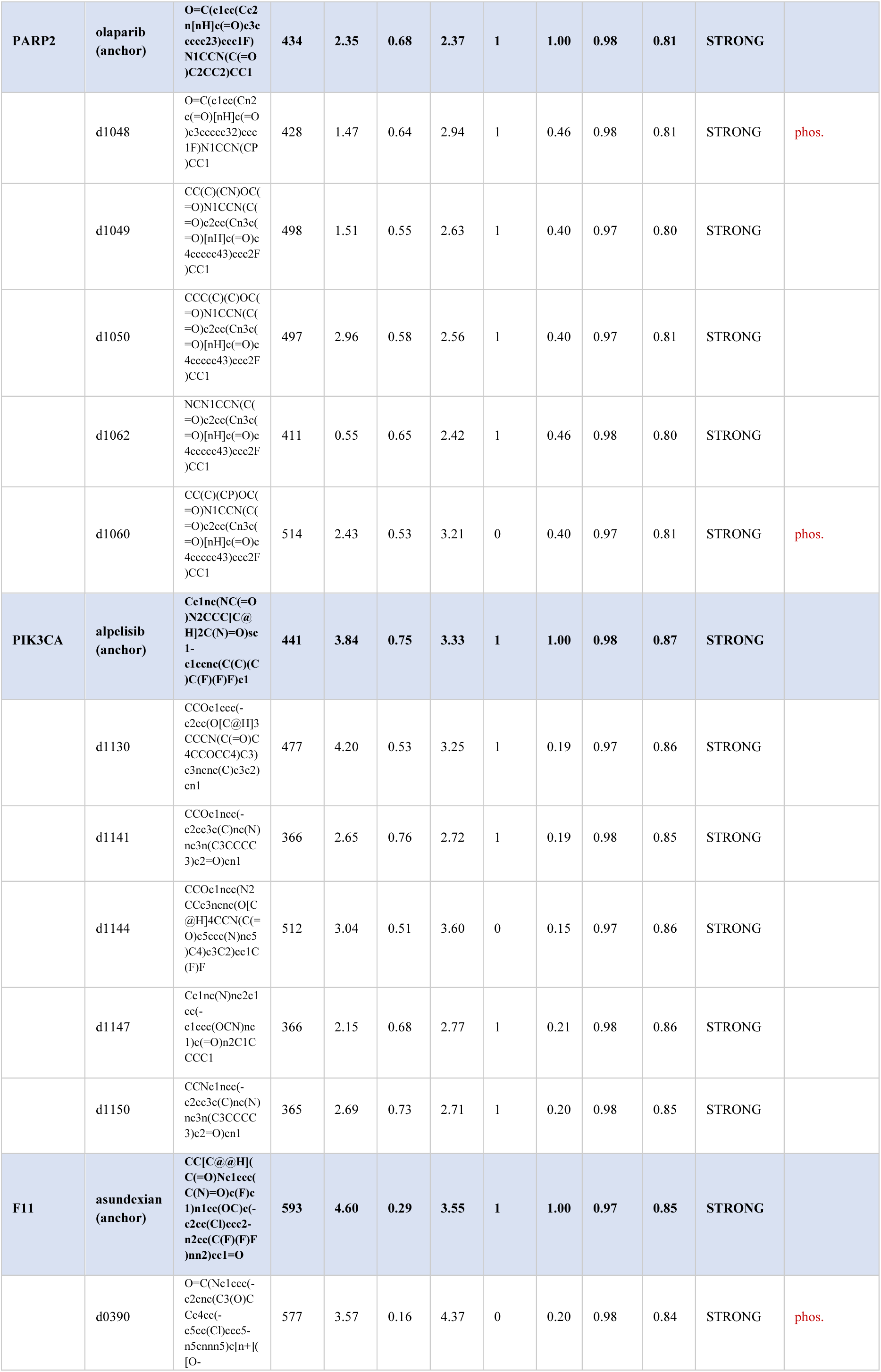
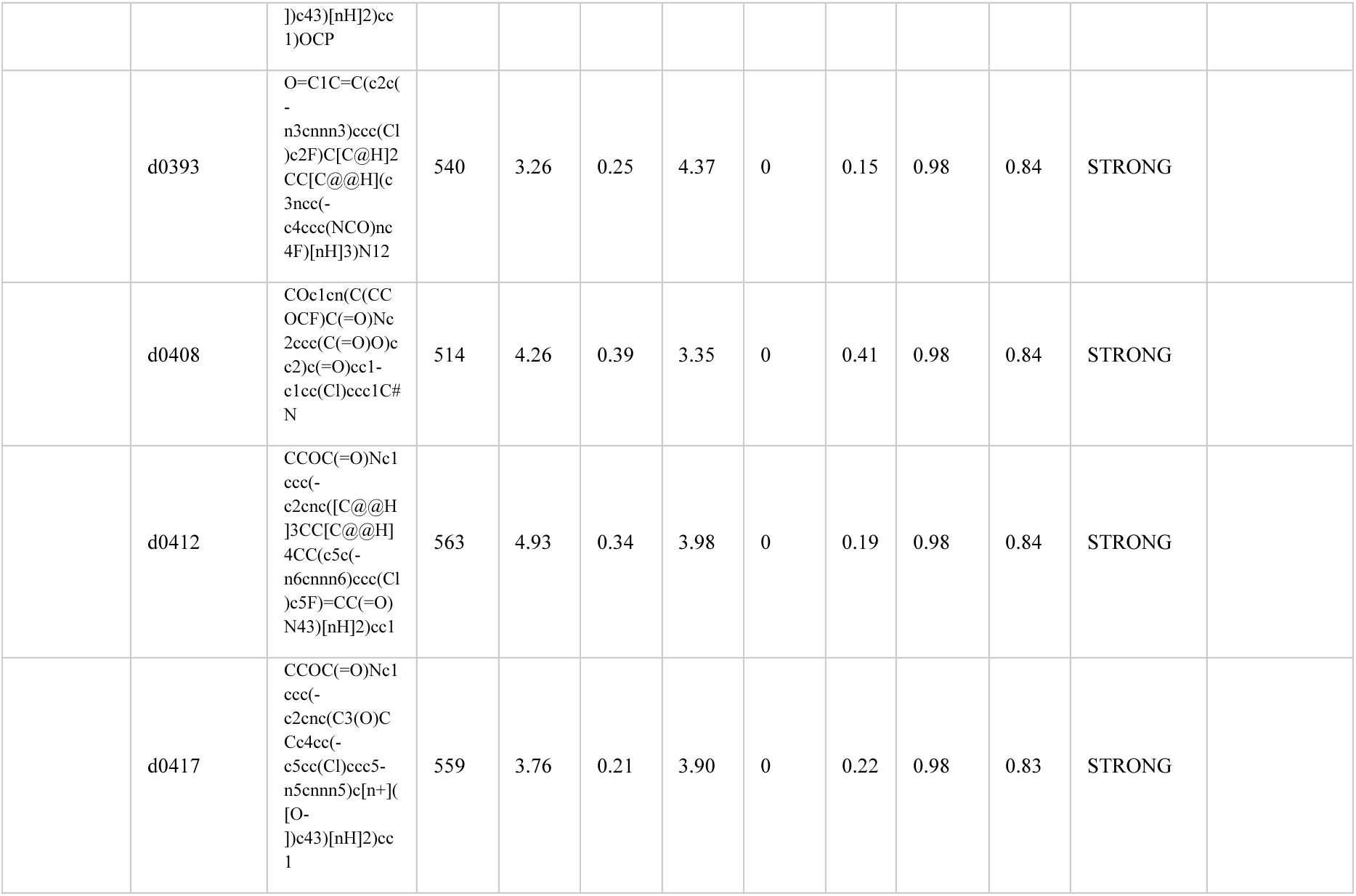
SMILES strings and computed properties of every dtSFM-generated design shown in the structural-validation gallery. (**Fig. 7).** The five designs per target — together with the approved-drug anchor for each of the 16 targets (shaded rows mark each target’s anchor and the start of a new target block). MW = molecular weight; QED = quantitative estimate of drug-likeness; SA = synthetic accessibility score (1 = easy, 10 = hard); Ro5 = Lipinski rule-of-five compliant (1/0); Tan = ECFP4 Tanimoto to the target’s approved drug; iPTM and PAE are AlphaFold 3 interface confidence metrics; AF3 = gate verdict (STRONG = iPTM ≥ 0.9 AND PAE ≤ 1.67 Å; MODERATE (MOD) = iPTM ≥ 0.7 AND PAE ≤ 5 Å; WEAK = below). ChemFlag marks designs carrying a structural alert — a reactive sp3 alkyl halide (alkyl-X), a bare phosphine not bonded to oxygen (phos.), or a free thiol (thiol); blank = none of these.

## References

1. Reddy, S. T. Computational convergence of adaptive immunity and artificial intelligence. bioRxiv (2026) doi:10.64898/2026.02.03.703525.

2. Reddy, S. T. Methods for molecular recognition computing. bioRxiv (2026) doi:10.64898/2026.04.03.716328.

3. Reddy, S. T. Vibe coding specificity foundation models. bioRxiv (2026) doi:10.64898/2026.06.04.730134.

4. Abramson, J. et al. Accurate structure prediction of biomolecular interactions with AlphaFold 3. Nature 630, 493–500 (2024).

5. Klaeger, S. et al. The target landscape of clinical kinase drugs. Science 358, eaan4368 (2017).

6. Macarron, R. et al. Impact of high-throughput screening in biomedical research. Nat. Rev. Drug Discov. (2011) doi:10.1038/nrd3368.

7. Davis, M. I. et al. Comprehensive analysis of kinase inhibitor selectivity. Nat. Biotechnol. 29, 1046–1051 (2011).

8. Renaud, J.-P. et al. Biophysics in drug discovery: impact, challenges and opportunities. Nat. Rev. Drug Discov. (2016) doi:10.1038/nrd.2016.123.

9. Erlanson, D. A., Fesik, S. W., Hubbard, R. E., Jahnke, W. & Jhoti, H. Twenty years on: the impact of fragments on drug discovery. Nat. Rev. Drug Discov. (2016) doi:10.1038/nrd.2016.109.

10. Hopkins, A. L., Keserü, G. M., Leeson, P. D., Rees, D. C. & Reynolds, C. H. The role of ligand efficiency metrics in drug discovery. Nat. Rev. Drug Discov. 13, (2014).

11. Wohlwend, J., et al. Boltz-1: Democratizing Biomolecular Interaction Modeling. bioRxiv (2024) doi:10.1101/2024.11.19.624167.

12. Passaro, S., et al. Boltz-2: Towards accurate and efficient binding affinity prediction. bioRxivorg (2025) doi:10.1101/2025.06.14.659707.

13. Watson, J. L. et al. De novo design of protein structure and function with RFdiffusion. Nature 620, 1089–1100 (2023).

14. Pacesa, M. et al. One-shot design of functional protein binders with BindCraft. Nature 646, 483–492 (2025).

15. Stark, H. et al. BoltzGen: Toward Universal Binder Design. bioRxiv (2025) doi:10.1101/2025.11.20.689494.

16. Schneuing, A. et al. Structure-based drug design with equivariant diffusion models (DiffSBDD). Nat. Comput. Sci. (2024) doi:10.1038/s43588-024-00737-x.

17. Guan, J. et al. 3D Equivariant Diffusion for Target-Aware Molecule Generation and Affinity Prediction. arXiv preprint arXiv:2303. 03543; ICLR 2023 (2023).

18. Peng, X., et al. Pocket2Mol: Efficient Molecular Sampling Based on 3D Protein Pockets. arXiv preprint arXiv:2205. 07249; ICML 2022 (2022).

19. Vaswani, A., et al. Attention is all you need. arXiv [cs.CL] (2017).

20. Luce, R. D. Individual Choice Behavior: A Theoretical Analysis. (Wiley, New Delhi, India, 1959).

21. Lee, H., et al. Contrastive learning for antibody-antigen sequence-to-specificity prediction. bioRxiv (2026) doi:10.64898/2026.02.25.707916.

22. Ross, J. et al. Large-scale chemical language representations capture molecular structure and properties. *Nat*. Mach. Intell. 4, 1256–1264 (2022).

23. Weininger, D. SMILES, a chemical language and information system. 1. Introduction to methodology and encoding rules. J. Chem. Inf. Comput. Sci. 28, (1988).

24. Lin, Z. et al. Evolutionary-scale prediction of atomic-level protein structure with a language model. Science 379, 1123–1130 (2023).

25. Liu, Z. et al. Forging the Basis for Developing Protein--Ligand Interaction Scoring Functions (PDBbind). Acc. Chem. Res. 50, (2017).

26. Lemos, B., et al. SAIR: Enabling Deep Learning for Protein-Ligand Interactions with a Synthetic Structural Dataset. bioRxiv (2025) doi:10.1101/2025.06.17.660168.

27. Mendez, D. et al. ChEMBL: towards direct deposition of bioassay data. Nucleic Acids Res. 47, (2019).

28. Gilson, M. K. et al. BindingDB in 2015: A public database for medicinal chemistry, computational chemistry and systems pharmacology. Nucleic Acids Res. 44, (2016).

29. Steinegger, M. & Söding, J. MMseqs2 enables sensitive protein sequence searching for the analysis of massive data sets. Nat. Biotechnol. (2017) doi:10.1038/nbt.3988.

30. Mangan, M. S. J. et al. Targeting the NLRP3 inflammasome in inflammatory diseases. Nat. Rev. Drug Discov. 17, 688 (2018).

31. Allard, B., Allard, D., Buisseret, L. & Stagg, J. The adenosine pathway in immuno-oncology. Nat. Rev. Clin. Oncol. 17, 611–629 (2020).

32. Decout, A., Katz, J. D., Venkatraman, S. & Ablasser, A. The cGAS-STING pathway as a therapeutic target in inflammatory diseases. Nat. Rev. Immunol. 21, 548–569 (2021).

33. Coll, R. C. et al. A small-molecule inhibitor of the NLRP3 inflammasome for the treatment of inflammatory diseases. Nat. Med. (2015) doi:10.1038/nm.3806.

34. Lawson, K. V. et al. Discovery of AB680: A Potent and Selective Inhibitor of CD73. J. Med. Chem. (2020) doi:10.1021/acs.jmedchem.0c00525.

35. Corrales, L. et al. Direct Activation of STING in the Tumor Microenvironment Leads to Potent and Systemic Tumor Regression and Immunity. Cell Rep. (2015) doi:10.1016/j.celrep.2015.04.031.

36. Madani, A. et al. Large language models generate functional protein sequences across diverse families. Nat. Biotechnol. 41, 1099–1106 (2023).

37. Bagal, V., Aggarwal, R., Vinod, P. K. & Priyakumar, U. D. MolGPT: Molecular Generation Using a Transformer-Decoder Model. J. Chem. Inf. Model. (2021) doi:10.1021/acs.jcim.1c00600.

38. Dauparas, J. et al. Robust deep learning--based protein sequence design using ProteinMPNN. Science (2022) doi:10.1126/science.add2187.

39. Cotet, T.-S., et al. Crowdsourced protein design: Lessons from the Adaptyv EGFR binder competition. bioRxiv (2025) doi:10.1101/2025.04.17.648362.

40. Sanjrani, N., Coupry, D. E., Pogány, P., Palmer, D. S. & Pickett, S. D. Benchmarking 3D structure-based molecule generators. J. Chem. Inf. Model. 65, 8006–8021 (2025).

41. van den Oord, A., Li, Y. & Vinyals, O. Representation Learning with Contrastive Predictive Coding (InfoNCE). arXiv preprint arXiv:1807. 03748 (2018).

42. Landrum, G. RDKit: Open-source cheminformatics software. doi:10.5281/zenodo.591637.

43. Loshchilov, I. & Hutter, F. Decoupled Weight Decay Regularization (AdamW). arXiv preprint arXiv:1711. 05101; ICLR 2019 (2019).

44. Mirdita, M. et al. ColabFold: making protein folding accessible to all. Nat. Methods 19, 679–682 (2022).

45. Rogers, D. & Hahn, M. Extended-Connectivity Fingerprints. J. Chem. Inf. Model. 50, (2010).

